# Chikungunya virus infection disrupts lymph node lymphatic endothelial cell composition and function via MARCO

**DOI:** 10.1101/2023.10.12.561615

**Authors:** Cormac J. Lucas, Ryan M. Sheridan, Glennys V. Reynoso, Bennett J. Davenport, Mary K McCarthy, Aspen Martin, Jay R. Hesselberth, Heather D. Hickman, Beth A. J. Tamburini, Thomas E. Morrison

## Abstract

Infection with chikungunya virus (CHIKV) causes disruption of draining lymph node (dLN) organization, including paracortical relocalization of B cells, loss of the B cell-T cell border, and lymphocyte depletion that is associated with infiltration of the LN with inflammatory myeloid cells. Here, we find that during the first 24 h of infection, CHIKV RNA accumulates in MARCO-expressing lymphatic endothelial cells (LECs) in both the floor and medullary LN sinuses. The accumulation of viral RNA in the LN was associated with a switch to an antiviral and inflammatory gene expression program across LN stromal cells, and this inflammatory response, including recruitment of myeloid cells to the LN, was accelerated by CHIKV-MARCO interactions. As CHIKV infection progressed, both floor and medullary LECs diminished in number, suggesting further functional impairment of the LN by infection. Consistent with this idea, we find that antigen acquisition by LECs, a key function of LN LECs during infection and immunization, was reduced during pathogenic CHIKV infection.

## INTRODUCTION

Chikungunya virus (CHIKV), a mosquito-transmitted arthritogenic alphavirus, remains a persistent threat to global health 10 years after spreading to the Americas and 20 years since epidemic-level outbreaks occurred in the Indian Ocean region (1–4). CHIKV disease typically presents with acute fever, rash, and severe arthralgia affecting the small joints, and up to 60% of patients remain chronically afflicted with arthritis and arthralgia for months to years post infection (2, 5). Chronic CHIKV musculoskeletal disease constitutes a high socioeconomic burden and warrants continued investigation of the mechanisms by which CHIKV infection causes chronic disease (5–9). Studies in both animal models (10–15) and human patients (16, 17) suggest that chronic CHIKV disease is associated with persistence of viral RNA and antigen in cells present in joint-associated tissue, such as synovial macrophages.

In prior studies using an immunocompetent mouse model of CHIKV infection, we found that CHIKV evades the B cell response to establish viral persistence in joint-associated tissues. These studies revealed that peripheral lymph nodes are important for the generation of the CHIKV-specific B cell response and control of CHIKV infection (12, 13, 18, 19). Indeed, the early appearance of CHIKV-specific IgG3 in human patients is associated with milder disease signs and symptoms and a shortened disease duration, emphasizing the importance of the antibody response for limiting acute and chronic CHIKV disease (20, 21). Closer examination of the lymph node response during CHIKV infection revealed that wild-type (WT) CHIKV infection disrupts the structure and function of the draining lymph node (dLN), the first secondary lymphoid organ to encounter virus following infection (22), in contrast to infection with the attenuated CHIKV 181/25 strain, which does not establish persistent infection in mice (18). The disruption of dLN organization during CHIKV infection is mediated by an early influx of inflammatory myeloid cells that impairs both lymphocyte recruitment and retention, loss of a clearly defined B-T cell border, and paracortical B cell re-localization, all of which contribute to reduced B cell germinal center formation (19, 22). However, the specific cell types that interact with CHIKV and promote inflammation in the dLN remain to be elucidated.

The architecture and cellular organization of LNs are essential for the development of effective immune responses against viral antigens (23). The organization of LNs is coordinated by lymph node stromal cells (LNSCs), including fibroblastic reticular cells (FRCs), lymphatic endothelial cells (LECs), and blood endothelial cells (BECs). These cell populations provide a physical scaffold for immune cell migration and produce signals to regulate migration, adhesion, localization, function, and survival of hematopoietic cells. FRCs and LECs can be grouped into distinct subsets based on transcriptional signatures and regional localization within the LN (24–29). LECs are among the first cells in the LN to encounter viruses, cells, and antigens draining into the LN via the afferent lymphatics (30–32). LN LECs specifically have been implicated in the maintenance of peripheral tolerance through AIRE-independent self-antigen presentation on MHC class II, which ultimately promotes the differentiation of FoxP3^+^ regulatory CD4^+^ T cells and limits activation of CD8^+^ T cells (33–37). More recently, LN LECs were shown to play an active role in enhancing immune responses through internalization and retention of antigen during vaccination and infection. This process, termed antigen archiving, supports protective memory T cell responses (38–40).

Recently, we found that CHIKV dissemination within an infected host is restricted by LNs and the scavenger receptor MARCO (macrophage receptor with collagenous structure), and that viral particles co-localize with MARCO^+^ LECs in the dLN following infection (41). Single cell mRNA sequencing (scRNA-seq) of LNSCs confirmed that viral RNA in the dLN accumulated largely in a subset of LECs that express MARCO termed MARCO^+^ LECs (41). As LECs are important regulators of LN tissue organization and function by establishing chemokine gradients that direct cell trafficking and by the secretion of cell maintenance factors, we hypothesized that the interaction between CHIKV and MARCO^+^ LECs promotes LN inflammation previously shown to impair B cell responses during CHIKV infection (42–48). In support of this idea, MARCO modulates the balance of protective and pathogenic inflammation in the lung to restrict acute influenza virus and pneumococcal lung infection and can enhance the inflammatory response to adenovirus infection (49–51), emphasizing that CHIKV-MARCO^+^ LEC interactions could promote LN inflammation and injury.

In this study, using scRNA-seq, we identified MARCO-expressing floor LECs that line the subcapsular sinus as a site of early viral RNA accumulation. These findings, together with our prior observations that CHIKV RNA subsequently accumulates in MARCO^+^ LECs that line the medullary sinuses, suggest that CHIKV targets multiple subsets of MARCO-expressing LECs in the LN. Flow cytometric and scRNA-seq analyses indicated that the number of LECs was not significantly altered early after infection. Instead, within the first 24 h, CHIKV infection caused dramatic alterations to the gene expression program of LNSCs, including LECs and other LNSC subtypes, that was characterized by an inflammatory gene transcriptional response, and this early inflammatory response was accelerated by CHIKV-MARCO interactions. However, quantification of LN LEC subsets throughout the acute phase of CHIKV infection revealed reduced numbers of both floor and medullary LECs at later times post-infection. Finally, evaluation of LN LEC function during CHIKV infection revealed that WT, but not attenuated, CHIKV infection impairs antigen acquisition by LN LECs. Collectively, these findings identify a role for the scavenger receptor MARCO in regulation of LN inflammation and support a model by which CHIKV targeting of MARCO-expressing LECs initiates an inflammatory response that rewires the transcriptional program of LNSCs, alters the composition of specialized LEC subtypes, and impairs known LEC functions.

## RESULTS

### CHIKV RNA accumulates in MARCO-expressing lymphatic endothelial cells in the dLN

Previously, using confocal microscopy and scRNA-seq analysis of dLN cells, we discovered that MARCO^+^ LECs lining the medullary sinuses internalize virus particles and harbor CHIKV RNA at 24 h post-infection (41). Notably, analysis of LEC subsets captured from mock- and CHIKV-infected LNs indicated that the number of floor LECs captured was reduced in CHIKV-infected compared with uninfected control LNs (41). Floor LECs line the inner layer of the subcapsular sinus (SCS) and have intimate contacts with antigen-detecting SCS macrophages (24, 43, 52), making them the first LNSCs to encounter cells and foreign antigens entering the LN and suggesting these cells could interact with CHIKV prior to MARCO^+^ LECs in the medullary sinus. CHIKV replication peaks between 24-72 h following infection of WT C57BL/6 mice (12, 13, 18, 41, 53), and can be directly cytopathic (54). Based on these observations, and previous reports that some floor LECs express MARCO (24, 25, 29), we hypothesized that floor LECs interact with CHIKV at earlier times post-infection leading to their alteration or reduction by 24 h post-infection. To test this idea, LNSCs from the dLN of mock- and CHIKV-infected mice were profiled by scRNA-seq at 8 h post-infection. Similar to our previous analysis at 24 h post-infection, LN cell suspensions were enriched for CD45^-^ cells by negative selection before processing for scRNA-seq (**Supplemental Figure 1**) (41). In comparison with our prior analysis of LNSCs at 24 h (41), fewer viral RNA counts were detected at 8 h. Thus, we enriched the cDNA libraries for CHIKV RNA using our previously described resampling and resequencing method (55). After enrichment, we classified cells with at least 1 viral RNA count as CHIKV^+^ and assessed the percentage of counts aligning to CHIKV for each individual cell (CHIKV score, **Figure 1A**). Analysis of cells harboring viral RNA revealed several CHIKV^+^ cell types mainly consisting of different endothelial cell and fibroblast populations (**Figure 1B**), a finding that is consistent with the known primary tropism of CHIKV for non-hematopoietic cells (15, 56–58). Among the CHIKV^+^ cell types identified, the populations with the highest CHIKV scores and the greatest proportion of CHIKV^+^ cells within that cell type were floor LECs and MARCO^+^ LECs (**Figure 1C and 1D**), with the floor LEC population containing the highest proportion of CHIKV^+^ cells.

**Fig 1.**
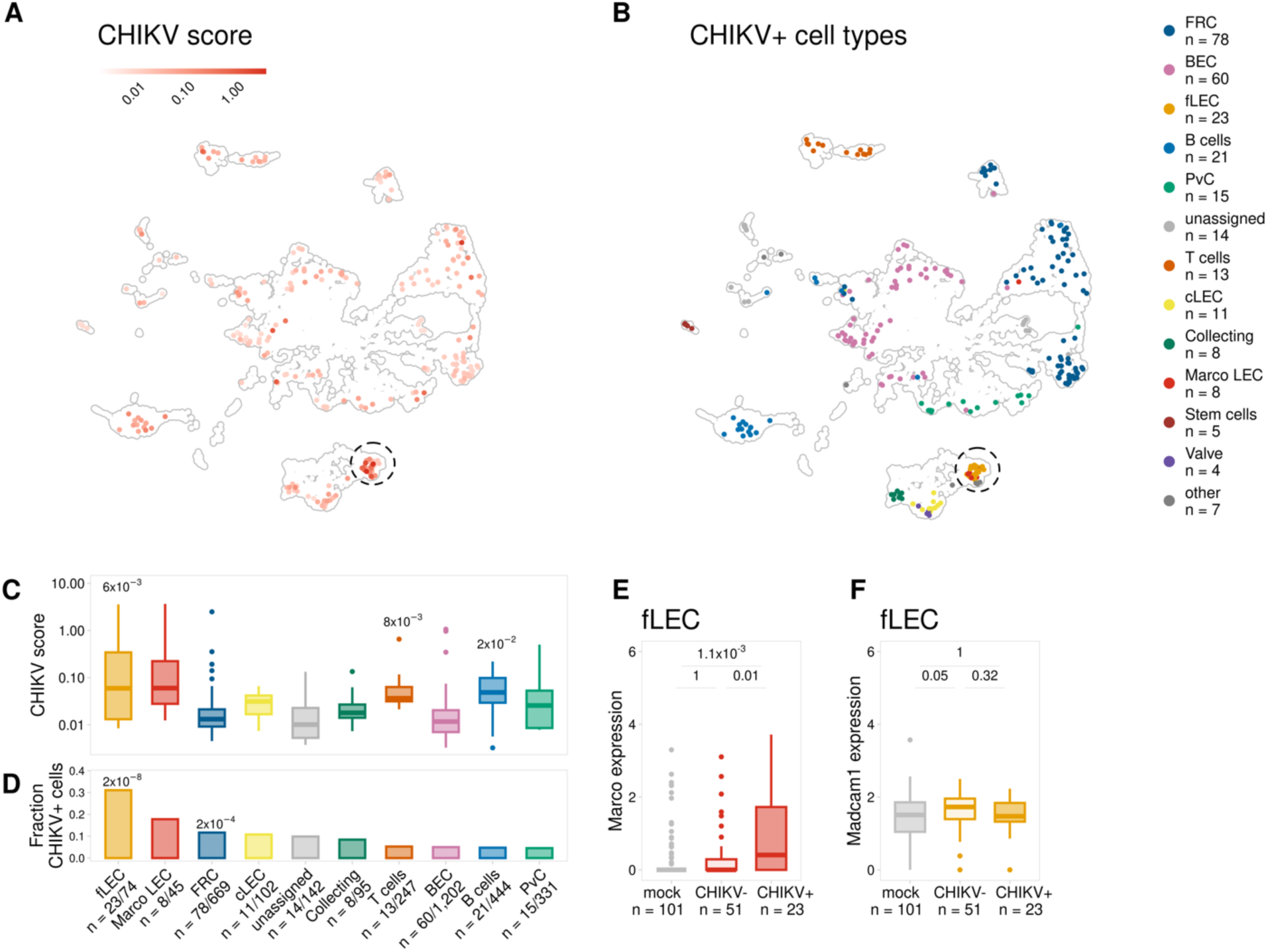
CHIKV RNA accumulates in MARCO-expressing floor LECs in the dLN. (**A-F**) WT C57BL/6 mice were inoculated with PBS (mock, n = 2) or 10^3^ PFU of CHIKV (n = 2) in the left-rear footpad. At 8 h post-infection, the dLN was collected and enzymatically digested into a single-cell suspension. Cells were enriched for CD45^-^ cells and analyzed by scRNA-seq as described in the Materials and Methods. (**A**) UMAP projection shows CHIKV score, calculated as the fraction of counts from the CHIKV-capture libraries aligning to the CHIKV genome. (**B**) UMAP projection shows CHIKV^+^ cells (>0 CHIKV counts). (**C**) CHIKV score is shown for CHIKV^+^ cells for cell types with >40 total cells and >3 CHIKV^+^ cells. P values were calculated using a one-sided Wilcoxon rank sum test with Bonferroni correction comparing each cell type with all other CHIKV^+^ cells. Only adjusted p-values <0.05 are shown. (**D**) The fraction of cells identified as CHIKV^+^ is shown for each cell type in C. P values were calculated using a one-sided hypergeometric test with Bonferroni correction. Labels show the number of CHIKV^+^ cells/total cells. Only adjusted p-values <0.05 are shown. (**E**) *Marco* and (**F**) *Madcam1* expression is shown for floor LECs (fLEC) for mock- and CHIKV-infected cells classified as either CHIKV^-^ or CHIKV^+^ (>0 CHIKV counts). *P* values were calculated using a two-sided Wilcoxon rank-sum test with Bonferroni correction. In the boxplots, the central lines, the box limits, and the whiskers represent medians, the interquartile range (IQR), and the min/max values that are not outliers, respectively. Outliers are shown as points and include any values that are more than 1.5x IQR away from the box.

Since CHIKV RNA was predominantly detected in MARCO-expressing LECs at 24 h post-infection (41), we next analyzed *Marco* expression in the CHIKV RNA^+^ and RNA^-^ floor LECs at 8 h post-infection. CHIKV RNA^+^ floor LECs (CHIKV^+^) exhibited greater *Marco* expression than CHIKV RNA^-^ floor LECs (CHIKV^-^) (**Figure 1E**). Floor and MARCO^+^ LECs are transcriptionally similar but can be distinguished by *Madcam1* expression in floor LECs, which is absent in MARCO^+^ LECs (24). In addition, floor LECs are characterized by a lower level and less uniform expression of *Marco* in contrast to MARCO^+^ LECs, which display a high level of MARCO expression (24). Both CHIKV RNA^+^ and CHIKV RNA^-^ floor LECs exhibited similar *Madcam1* expression, a marker unique to floor LECs (59–61), suggesting that these cells are indeed floor LECs and not mis-annotated MARCO^+^ LECs (**Figure 1F**). Collectively, these studies reveal that CHIKV RNA accumulates in two subsets of LECs during the first 24 h of infection and suggest that CHIKV interactions with MARCO are important for viral capture and internalization by endothelial cells in the LN.

### CHIKV RNA^+^ LECs show signs of active CHIKV replication

At 24 h, CHIKV-high cells identified by scRNA-seq had attributes consistent with decreased cell viability and with virus-mediated transcriptional shutoff, including expression of fewer host genes and an increased fraction of reads aligning to mitochondrial genes (41), suggesting that LECs support active CHIKV replication. One marker of active CHIKV RNA replication is the production of a positive-sense subgenomic mRNA that encodes the viral structural polyprotein (62). To look for further evidence of viral replication, we calculated the ratio (sgRNA ratio) of reads aligning to the viral sgRNA and reads aligning upstream of the sgRNA, which should reflect levels of the full-length viral genome. Consistent with the localization of CHIKV-high cells in our previous study (41), at 24 h cells with the highest sgRNA ratio were found predominantly within the MARCO^+^ LEC subset and a cluster of endothelial cells that we were unable to further annotate due to the low number of expressed host genes (unassigned-LEC)

(**Supplemental Figure 2A-D).** When we further characterized cell types with the highest CHIKV sgRNA ratio at 24 h, we observed a negative correlation between the sgRNA ratio and the number of mouse genes expressed by MARCO^+^ LECs and unassigned-LECs (**Supplemental Figure 2E**). In addition, we also identified a positive correlation between the sgRNA ratio and the percentage of mitochondrial reads per cell (**Supplemental Figure 2E**). Notably, the unassigned-LECs have both a higher CHIKV sgRNA ratio and percentage of mitochondrial reads and a lower number of expressed mouse genes compared with MARCO^+^ LECs, suggesting these cells could be severely injured MARCO^+^ LECs (**Supplemental Figure 2E**). Overall, these results show that cells with high levels of viral sgRNA also have indications of reduced viability, suggesting that CHIKV replication occurs in LECs that capture and internalize CHIKV particles and leads to cell injury or death.

### Lymph node sinus alteration during pathogenic CHIKV infection

Our analyses indicated that CHIKV RNA accumulates in multiple subsets of MARCO-expressing LN LECs during infection. To investigate the fate of these cells, we evaluated Lyve-1 and MARCO expression in the dLN during infection with the attenuated CHIKV 181/25 strain, which does not disrupt dLN cellular organization (22), and the parental WT CHIKV AF15561 strain at 8, 24, and 48 h post-infection using immunofluorescent confocal microscopy. At 8 h post-infection, Lyve-1 and MARCO expression were similar across dLNs from mock-, CHIKV 181/25-, and WT CHIKV-infected mice (**Supplemental Figure 4A**). Lyve-1 signal was observed in both subcapsular and medullary sinus regions, supporting the annotation of both floor and MARCO^+^ LEC subsets in the 8 h post-infection scRNA-seq dataset and consistent with previous reports indicating specificity of Lyve-1 expression for floor (low level), MARCO^+^ (high level), and Ptx3 (high level) LECs (24, 25, 63). MARCO signal was localized predominantly to the LN medullary sinuses, consistent with reports indicating MARCO expression on LN LECs is regionally distinct (24, 25). At 24 h post-infection, Lyve-1 and MARCO expression remained similar in dLNs from mock-, CHIKV 181/25-, and WT CHIKV-infected mice (**Supplemental Figure 4B**). However, by 48 h post-infection, Lyve-1 signal was greatly reduced and MARCO signal was undetectable in dLNs from WT CHIKV-infected mice (**Figure 2A** and **2B** and **Supplemental Figure 4C**). Higher magnification imaging of the subcapsular and medullary sinuses of LNs from WT CHIKV-infected mice revealed marked expansion of the LN sinuses (**Figure 2C**). Lyve-1^+^ bleb-like structures were visible within the expanded sinus, suggesting the presence of damaged cells (**Figure 2C**). To investigate the cellular composition of the expanded sinuses, LN sections from WT CHIKV-infected mice were stained for CD11b, as prior studies had identified localization of inflammatory monocytes to the medullary region of the dLN at 24 h post-infection (19), and DAPI to identify cell nuclei. Indeed, the expanded LN sinuses contained numerous CD11b^+^ cells (**Figure 2D**), supporting the association of inflammatory cellular infiltrates with alteration of resident cells in the LN sinus.

**Figure 2.**
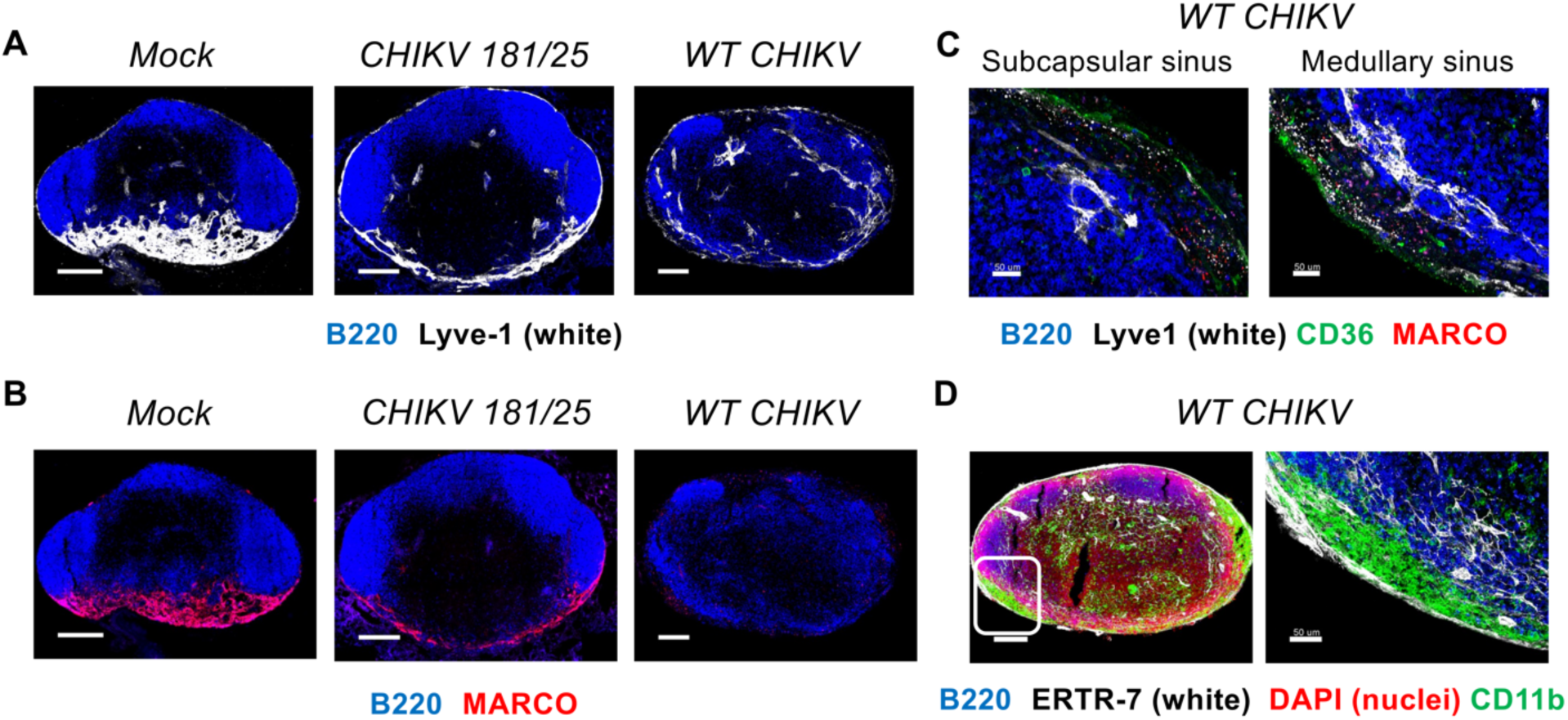
WT CHIKV infection disrupts LEC marker expression and elicits infiltration of LN sinuses. (**A-D**) WT C57BL/6 mice were mock-inoculated (n = 3) or inoculated in the footpad with 10^3^ PFU CHIKV 181/25 (n = 5) or WT CHIKV (n = 5) and the dLN was collected at 48 h post-infection. (**A**) Frozen dLN sections were stained for B220 (B cells, blue) and Lyve-1 (LECs, white). Scale bar, 200 μm. (**B**) Frozen dLN sections were stained for B220 (B cells, blue) and MARCO (red). Scale bar, 200 μm. (**C**) Higher magnification images of subcapsular and medullary sinus regions in LNs from WT CHIKV-infected mice stained for B220 (B cells, blue), Lyve-1 (floor and medullary LECs, white), CD36 (ceiling LECs, green), and MARCO (red). Scale bar, 50 μm. (**D**) LNs from WT CHIKV-infected mice stained for B220 (B cells, blue), ERTR-7 (fibroblasts, white), nuclei (red), and CD11b (myeloid cells, green). Scale bar, 200 μm (left), 50 μm (right). Images are representative of 3-5 dLNs per group (2 independent experiments).

### Pathogenic CHIKV infection alters LN LEC composition

To evaluate whether loss of Lyve-1 and MARCO signal at 48 h post-infection corresponded to the loss of specific LECs or an altered composition of LEC subsets, dLN stromal cells were evaluated at 1, 2, and 5 d post-infection by flow cytometry. After excluding non-viable cells, LNSCs were segregated within all CD45^-^ cells based on expression of CD31/PECAM-1 and podoplanin (PDPN) (24, 59, 60): CD31^+^PDPN^-^ blood endothelial cells (BECs), CD31^-^PDPN^+^ fibroblastic reticular cells (FRCs), and CD31^+^PDPN^+^ LECs. Lyve-1 and MARCO antibodies exhibit low, weak signal with flow cytometry (41, 64), thus, LECs subsets were defined using mannose receptor C-type 1 (MRC1), intercellular adhesion molecule 1 (ICAM1), integrin subunit alpha 2B (ITGA2B), and the scavenger receptor CD36, which have been used successfully in other studies to discriminate LEC subsets (25, 59, 64) including medullary LECs (MRC1^+^ICAM1^+^), floor LECs (MRC1^-^ICAM1^+^ITGA2B^+^), and ceiling LECs (MRC1^-^ICAM1^-^CD36^+^) (**Figure 3A**). The total number of BECs, FRCs, and LECs was similar between mock-, CHIKV 181/25-, and WT CHIKV-infected LNs at 1 and 2 d post-infection (**Figure 3B**). At 5 d post-infection, while the number of BECs increased to a similar extent following infection with either CHIKV 181/25 or WT CHIKV (**Figure 3B**), FRC and LEC numbers increased in CHIKV 181/25-infected LNs but not WT CHIKV-infected LNs (**Figure 3B**), suggesting that WT CHIKV infection alters the proliferation or survival of these LNSC subtypes. Further segregation of LECs into medullary, floor, and ceiling subtypes revealed that both the percentage and number of medullary LECs was reduced during WT CHIKV infection compared with CHIKV 181/25 infection (44.4-fold decrease in number, *P* < 0.0001), whereas the number of ceiling LECs was unchanged at 5 d post-infection (**Figure 3C**). The percentage and number of floor LECs was also reduced during WT CHIKV infection compared with CHIKV 181/25 infection (6.3-fold decrease in number, *P* < 0.0001) (**Figure 3C**). Notably, there was a small but significant reduction in the percentage and number (3-fold, *P* < 0.0044) of dLN floor LECs between mock and WT CHIKV-infected mice at 1 d post-infection, suggesting that these cells, which interact with the virus early after infection, could be damaged as a result. These data suggest that WT CHIKV infection impairs the expansion and/or maintenance of specific regional LEC subsets. Notably, medullary and floor LECs include all the MARCO-expressing LECs in the LN, which are LN cell types predominantly targeted by CHIKV early after infection (**Figure 1B-D**) (41), suggesting that these changes could be due to CHIKV-MARCO interactions.

**Figure 3.**
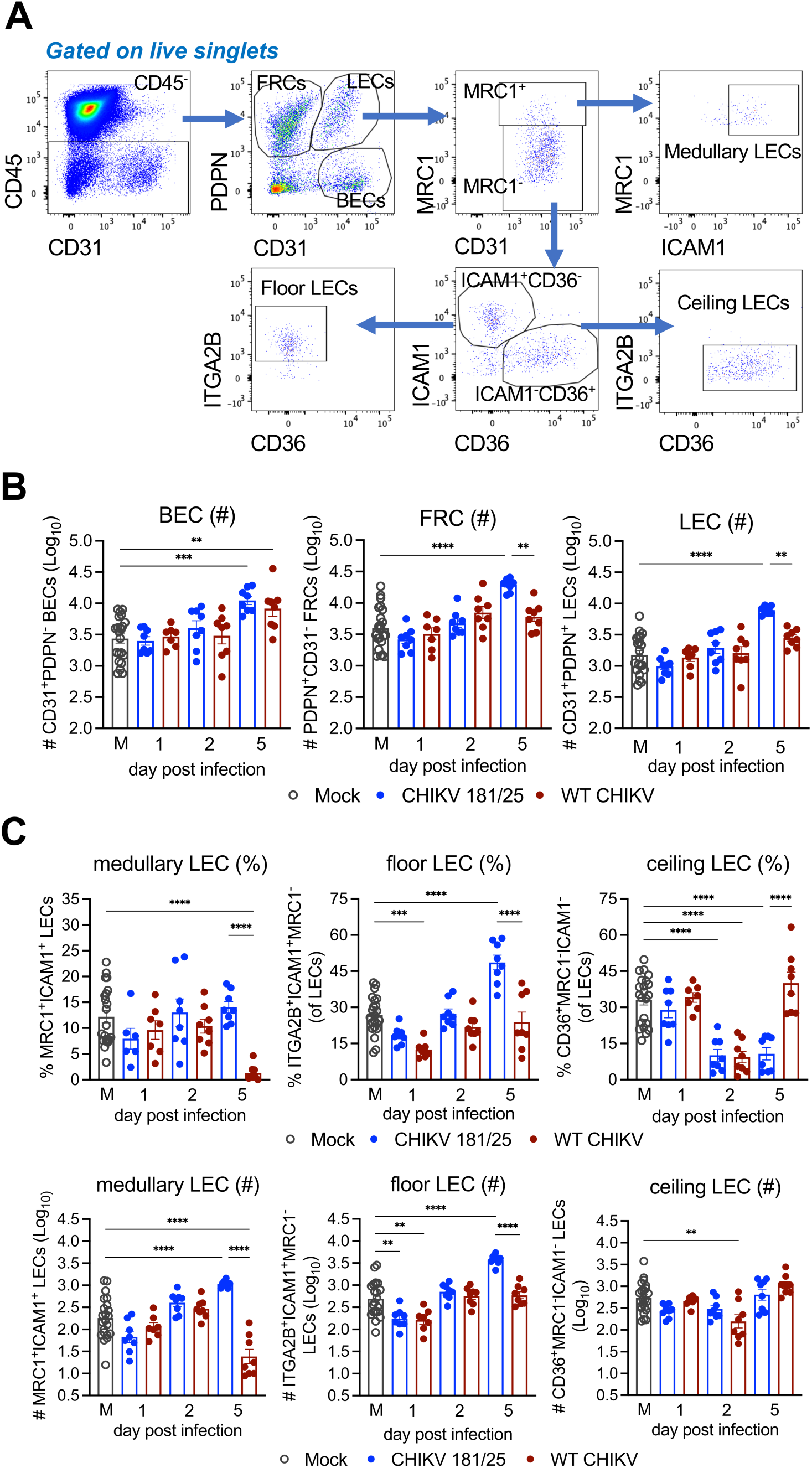
WT CHIKV infection alters LN LEC composition. (**A-C**) WT C57BL/6 mice were inoculated in the footpad with PBS (n = 8) or 10^3^ PFU CHIKV 181/25 (n = 8) or WT CHIKV (n = 8). At the timepoints indicated, the dLN was collected and analyzed by flow cytometry. (**A**) Gating strategy (after gating on viable singlets) to segregate BECs, LECs, and FRCs, and further subset LECs into floor, ceiling, and medullary LECs. (**B**) Total number of BECs, FRCs, and LECs in the dLN at each timepoint. (**C**) Percentage and total number of medullary, floor, and ceiling LECs in the dLN at each timepoint. Data are combined from 2 independent experiments and presented as mean +/-SEM. *, *P* < 0.05; **, *P* < 0.01; ***, *P* < 0.001; ****, *P* < 0.0001, one-way ANOVA with Tukey’s multiple comparison test.

### Alteration of LN LEC composition is dependent on CHIKV-MARCO interactions

Our prior studies identified a role for the scavenger receptor MARCO in early viral accumulation in the dLN and restricting early viral dissemination to distal tissues (41, 65). Furthermore, our data indicates that CHIKV RNA accumulates predominantly in subsets of LECs that express MARCO. Thus, we hypothesized that CHIKV-MARCO interactions promote the diminished Lyve-1 expression and the altered sinuses observed in the dLN of WT CHIKV-infected mice. To assess this, dLNs were evaluated at 48 h post-infection by confocal microscopy following CHIKV infection of WT and *MARCO^-/-^* mice. In contrast to the sparse Lyve-1 signal in the dLN of WT mice, the dLN of CHIKV-infected *MARCO^-/-^* mice exhibited more robust Lyve-1 expression (**Figure 4A).** Higher magnification imaging of the medullary sinus highlights the difference in Lyve-1 expression in the dLN of CHIKV-infected WT and *MARCO^-/^*^-^ mice, where Lyve-1 signal is substantially diminished in WT mice while *MARCO^-/-^*mice maintain robust Lyve-1 expression (**Figure 4B**). These data suggest that MARCO promotes the loss of Lyve-1 expression or LECs in the dLN during CHIKV infection. Furthermore, analysis of LN LEC subsets at 5 d post-infection revealed that in WT CHIKV-infected *MARCO^-/-^* mice, medullary and floor LEC populations were increased compared with WT CHIKV-infected WT mice (14.62-fold, *P* < 0.0001; 3.78-fold, *P* = 0.0214, increase in number respectively) (**Figure 4C-D**), indicating that the altered composition of LECs during CHIKV infection is MARCO-dependent. In addition, the percentage of ceiling LECs was reduced in WT CHIKV-infected *MARCO^-/-^*mice to a level comparable to CHIKV 181/25-infected WT mice in contrast to the higher percentage of ceiling LECs in WT CHIKV-infected WT mice (**Figure 4C**), suggesting that the change in the proportion of ceiling LECs may be a component of the LNSC response to pathogenic CHIKV infection. Overall, these data demonstrate a role for MARCO in the alteration of specific LN LEC populations during pathogenic WT CHIKV infection.

**Figure 4.**
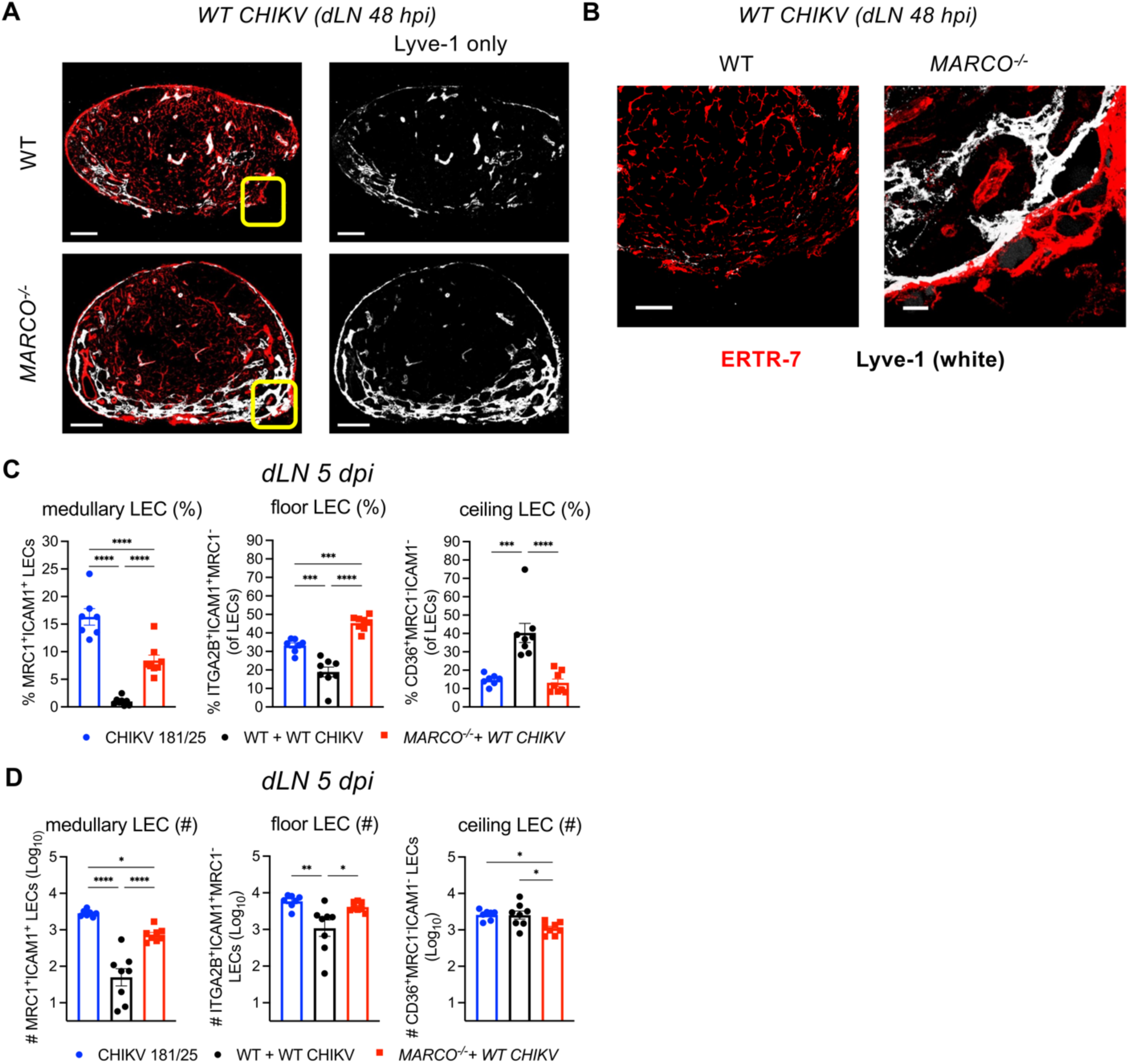
Loss of Lyve-1 and changes in LEC subset composition are MARCO-dependent. (**A-B**) WT and *MARCO^-/-^* C57BL/6 mice were infected in the footpad with 10^3^ PFU WT CHIKV (n = 7-8). At the indicated timepoints, the dLN was collected for analysis by immunofluorescent confocal microscopy or flow cytometry. (**A**) Frozen dLN sections were stained for Lyve-1 (LECs, white) and ERTR-7 (fibroblasts, red). Scale bar, 200 μm. (**B**) Higher magnification images of the indicated LN sinus region (yellow box). Scale bar 50 μm. Images are representative of 5 dLNs per group (2 independent experiments). (**C**) Percentage and (**D**) total number of medullary, floor, and ceiling LECs in the dLN at 5 d post-infection. Data are combined from 2 independent experiments and presented as mean +/-SEM. *, *P* < 0.05; **, *P* < 0.01; ***, *P* < 0.001; ****, *P* < 0.0001, one-way ANOVA with Tukey’s multiple comparison test.

### Inflammatory gene expression in LNSCs during CHIKV infection

We next evaluated differential gene expression in LNSCs from mock and CHIKV-infected mice at 8 and 24 h post-infection using our scRNA-seq datasets. UMAP projection of all LNSCs from mock- and CHIKV-infected mice at both 8 and 24 h post-infection revealed that LNSCs from CHIKV-infected mice at 8 h clustered strongly with LNSCs from mock-infected mice whereas LNSCs from CHIKV-infected mice at 24 h were strongly segregated from the other populations (**Figure 5A and 5B**) and this was consistent when coloring the UMAP projection by cell type (**Figure 5B**). We next identified gene ontology terms (biological process) for genes upregulated in each cell type. To identify the predominant gene expression programs upregulated during the first 24 h of CHIKV infection, we clustered gene ontology (GO) terms for each timepoint into distinct modules based on similarity. From this analysis the primary gene expression module upregulated at 8 h post infection consists of factors associated with the innate immune response (**Figure 5C**), including *Bst2* which is broadly upregulated in most cell types at 8 h and has been previously shown to promote retention of CHIKV particles at the host cell membrane to prevent virus release (66, 67). We also detected upregulation of *Zbp1*, a key factor in sensing cytosolic DNA during virus-induced cell damage and promoting production of anti-viral cytokines, and *Irf7*, another key factor in the induction of anti-viral cytokines such as interferon-β (68, 69) (**Figure 5D**). When we compare changes in gene expression between the 24 h and 8 h timepoints we identify a similar innate immune response module and observe further upregulation of *Bst2*, *Zbp1*, and *Irf7* (**Figure 5D-E**). By 24 h post-infection, we also detected increased expression of genes involved in a broader inflammatory response, including *Ccl2*, *Cxcl9*, *Cxcl10*, and *Ccl7*, which were upregulated across the major LNSC subsets (**Figure 5E-F**). However, minor differences in the degree of expression of specific genes were noted, with *Ccl7* upregulated to a lower degree in MARCO^+^ LECs and perivascular cells (PvCs) than FRCs, which is consistent with existing knowledge of fibroblasts being one of the primary CCL7-producing stromal cell types (**Figure 5F**) (70, 71). In addition, CCL2 is a potent chemoattractant for monocytes, which previous studies demonstrated are detrimental to LN structure and downstream anti-CHIKV antibody responses (19, 72); the high expression of *Ccl2* detected in LECs suggests these cells may contribute to early recruitment of inflammatory monocytes. We also evaluated expression of important LNSC homeostatic chemokines including *Ccl21a*, *Il7*, *Cxcl13*, and *Ccl19* (**Figure 5G**). The expression level of these genes was largely unchanged or diminished in LNSCs of CHIKV-infected mice when compared to mock-infected mice (**Figure 5G**), indicating that the primary effect of CHIKV infection on the transcriptome of LNSCs is activation of antiviral and inflammatory gene expression programs.

**Figure 5.**
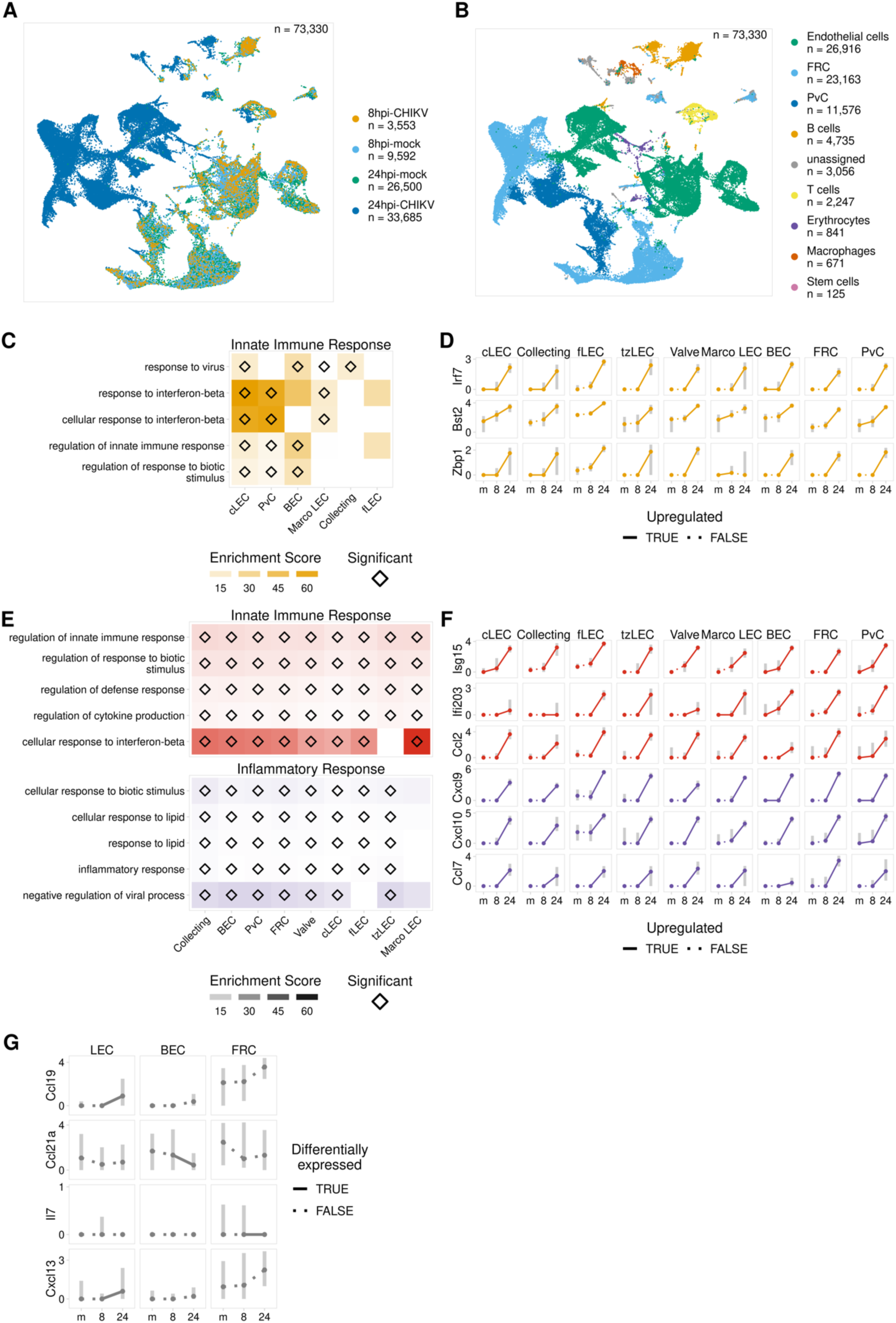
LNSCs exhibit a dominant pro-inflammatory response after 8 h post CHIKV infection. (**A-G**) Cells from the dLN were collected from mock- or WT CHIKV-inoculated mice at 8 or 24 h post-infection. Cells were subjected to CD45 depletion and then scRNA-seq. (**A**) UMAP projection showing all cells from the 8 and 24 h timepoints colored by sample. (**B**) UMAP projection showing all cells from the 8 and 24 h timepoints colored by cell type. (**C**) Enrichment scores for each cell type for the top 5 terms from the primary gene ontology module identified for the 8 h timepoint. Enrichment score is the fraction of upregulated genes overlapping the term divided by the fraction of background genes overlapping the term. Significantly enriched GO terms are marked by a diamond. (**D**) A selection of top upregulated genes for terms significantly enriched at 8 h. Points show the median expression for mock (m), 8 h (8), and 24 h (24) samples; grey bars show the interquartile range. A solid line indicates the gene is significantly upregulated between the timepoints. (**E**) Enrichment scores for the primary GO modules for the 24 h timepoint, as described in (C). (**F**) A selection of top upregulated genes for the terms identified at 24 h, plotted as described in (D). (**G**) Expression of select homeostatic chemokines among the major LNSC types, plotted as described in (D). A solid line indicates the gene is differentially expressed between the timepoints.

### MARCO expression triggers a rapid LN inflammatory response

The presence of inflammatory CD11b^+^ cells in the expanded sinuses of CHIKV-infected LNs (**Figure 3B** and **3C**), retention of Lyve-1 signal in *MARCO^-/-^* mice (**Figure 4A**), and high expression of monocyte chemoattractant *Ccl2* in LNSCs at 24 h post-infection (**Figure 5F**) suggest that CHIKV-MARCO interactions promote LN inflammation via regulation of inflammatory chemokine expression and recruitment of inflammatory monocytes. To investigate this hypothesis, inflammatory chemokine mRNA expression (*Ccl2*, *Cxcl1*, *Cxcl9*, and *Cxcl10*) was assessed in whole LNs from WT and *MARCO^-/-^* mice at time points (8, 12, and 16 h post-infection) prior to the dominant type I IFN response observed at 24 h (**Figure 5**) (19). Consistent with our scRNA-seq analysis, little to no upregulation of these chemokines was observed in CHIKV-infected WT or *MARCO^-/-^*mice at 8 h post-infection. However, by 12 h post-infection, the expression of *Ccl2*, *Cxcl1*, *Cxcl9*, and *Cxcl10* in the dLN of CHIKV-infected WT mice, but not *MARCO^-/-^* mice, was increased in comparison with mock-infected mice. Indeed, expression of *Ccl2*, *Cxcl1*, *Cxcl9*, and *Cxcl10* in the dLN of CHIKV-infected WT mice was increased 46.9-fold, 54.2-fold, 16.2-fold, and 17.2-fold over the expression level observed in *MARCO^-/-^* mice (**Figure 6A**). By 16 h post-infection, chemokine expression was increased in both CHIKV-infected WT and *MARCO^-/-^* mice in comparison with mock-infected mice, however, *Ccl2*, *Cxcl1*, and *Cxcl9* expression remained significantly higher in the dLN of WT mice (2.4-fold, 2.9-fold and 2.1-fold, respectively) (**Figure 6A**). To further address whether direct CHIKV-MARCO interactions and subsequent viral internalization promote early inflammatory chemokine expression in the dLN, chemokine expression in the dLN was assessed at 12 h post-infection in WT mice infected with WT CHIKV or CHIKV^E2^ ^K200R^, which lacks interaction with MARCO (41, 65, 73). The expression of *Ccl2* and *Cxcl1*, was increased 4-fold and 10.1-fold, respectively in the dLN of WT CHIKV-infected mice compared with CHIKV^E2^ ^K200R^-infected mice (**Figure 6B**), whereas differences in *Cxcl9* and *Cxcl10* expression were not statistically significant. These data suggest that, at least partially,

**Figure 6.**
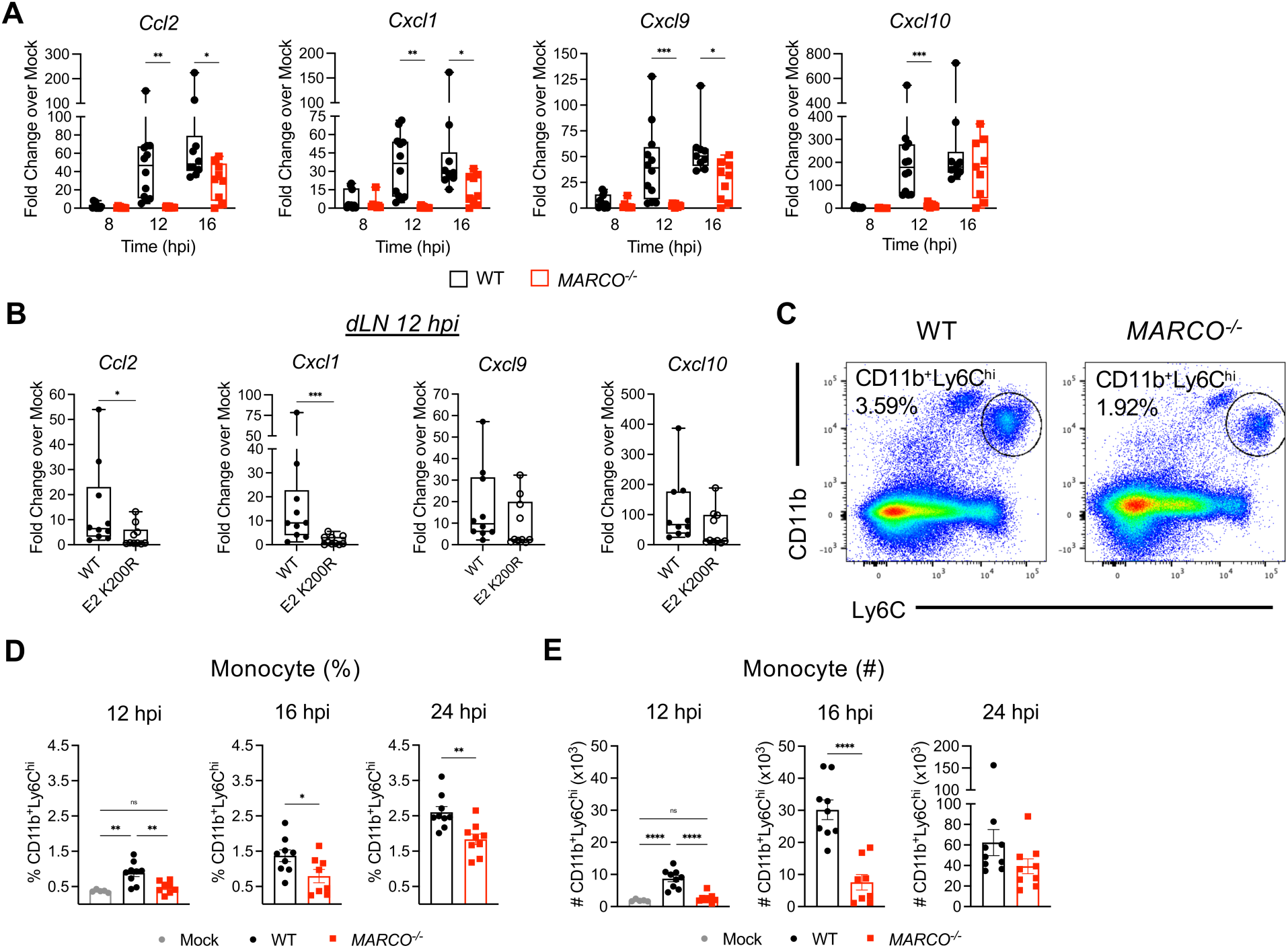
CHIKV-MARCO interactions promote LN inflammation. (**A-E**) WT and *MARCO^-/-^*C57BL/6 mice were inoculated in the footpad with 10^3^ PFU WT CHIKV (n = 9-13) or CHIKV^E2^ ^K200R^ (n = 10). At the indicated timepoints, the dLN was collected for gene expression analysis by RT-qPCR (**A-B**) or for infiltration of inflammatory myeloid cells by flow cytometry (**C-E**). (**A**) Expression of inflammatory chemokines in the dLN of CHIKV-infected WT and *MARCO^-/-^*mice at 8, 12, and 16 h post-infection. (**B**) Expression of inflammatory chemokines in the dLN of WT mice inoculated with WT CHIKV or CHIKV^E2^ ^K200R^ at 12 h post-infection. (**C**) Representative flow cytometry plots of inflammatory CD11b^+^Ly6C^hi^ monocytes. (**D**) Percentage of monocytes in the dLN at 12, 16, and 24 h post-infection. (**E**) Total number of monocytes per dLN at 12, 16, and 24 h post-infection. Data representative of 2-3 independent experiments. Statistical significance was determined by one-way ANOVA with Tukey’s multiple comparison test (**A**) or student’s t-test (**B**, **D**, **E**). *, *P* <0.05; **, *P* < 0.01; ***, *P* < 0.001; ****, *P* < 0.0001.

### CHIKV-MARCO interactions and viral internalization promote inflammatory gene expression in the dLN

To determine if MARCO also promotes infiltration of the LN with CD11b^+^ cells during CHIKV infection (**Figure 3**), we evaluated accumulation of inflammatory monocytes in the dLN of WT and *MARCO^-/-^* mice using flow cytometry. Consistent with higher *Ccl2* expression in WT mice at 12 and 16 h post-infection, the percentage (**Figure 6C**) and total number of (**Figure 6D**) inflammatory Ly6C^hi^ monocytes in the dLN (CD45^+^CD11c^neg^CD11b^+^Ly6C^hi^Ly6G^neg^) was significantly greater in WT mice at 12 (3.3-fold, *P* < 0.0001) and 16 h post-infection (4-fold, *P* < 0.0001) compared with *MARCO^-/-^*mice. By 24 h post-infection, the percentage of inflammatory monocytes remained significantly higher in WT mice than in *MARCO^-/-^* mice (**Figure 6C**) although the difference in monocyte numbers was not statistically significant (**Figure 6D**). These data suggest CHIKV-MARCO interaction induces a rapid early pro-inflammatory response, recruiting pathogenic monocytes which cause disruption of dLN cellular organization and impair B cell responses (19).

### Pathogenic, but not attenuated, CHIKV infection impairs foreign antigen acquisition by LECs

A key function of LN LECs is the ability to acquire and retain foreign antigen to promote long-lived adaptive immunity following both viral infection and vaccination (30, 38–40). Indeed, prior studies showed that fluorescently-labeled ovalbumin (ova) or influenza nucleoprotein (NP) is specifically acquired by LN LECs upon subcutaneous injection of mice experiencing an active viral infection or when delivered with an adjuvant such as polyI:C (38–40). To evaluate the functional capacity of LN LECs to acquire antigen during CHIKV infection, we immunized CHIKV 181/25- or WT CHIKV-infected mice with 10 µg ova-488 in both calf muscles (20 µg/mouse total) at 3 d post-infection, at which point the dLN of WT CHIKV-infected mice is substantially disorganized (22), and assessed the proportion of ova^+^ LNSCs 2 and 7 days later (**Figure 7A**). As a positive control, we immunized naïve mice in both calf muscles with 10 µg ova-488 and 5 µg polyI:C per calf (38, 39). Consistent with prior studies (38, 39), ova acquisition was highly specific to LN LECs, as little to no ova was detected in BECs or FRCs (**Supplemental Figure 5A**), confirming that we could measure LEC antigen acquisition by this method (39). We evaluated ova^+^ LNSCs in both the popliteal (first footpad draining LN) and iliac (next draining LN in sequence) LNs (74) to determine if any differences in ova acquisition were associated with impaired lymphatic drainage. Comparison of LNSC populations in the popliteal LN of ova-immunized/CHIKV-infected mice revealed that the number of LECs in WT CHIKV-infected mice was diminished at both 2 (7.3-fold, *P* < 0.0001) and 7 days (3.41-fold, *P* < 0.0021) post ova injection in the popliteal LN compared with CHIKV 181/25-infected mice (**Figure 7B**). FRC numbers were slightly decreased in WT CHIKV-infected mice compared with CHIKV 181/25 infection at both 2 and 7 days post ova immunization (2.42-fold, *P* < 0.1067; 2.7-fold, *P* < 0.0528, respectively) in the popliteal LN (**Figure 7B**), however, these differences were not statistically significant, and BEC numbers remained similar across all groups and time post immunization (**Figure 7B**), suggesting the adverse effects of WT CHIKV are specific to LECs. After gating on ova^+^ LECs in naïve mice as a negative control, we found that both the percentage and number of ova^+^ LECs was reduced during WT CHIKV-infection compared with CHIKV 181/25 infection in the popliteal LN (**Figure 7C-D**). The difference in ova^+^ LEC number was greater at both 2 and 7 days post-ova in the popliteal LN between CHIKV 181/25 and WT CHIKV infection (14-fold, *P* < 0.0001; 29.5-fold, *P* < 0.0001, respectively) than total LEC number alone (7.3-fold, *P* < 0.0001; 3.41-fold, *P* < 0.0021), suggesting differences in ova acquisition within the popliteal LN cannot be attributed solely to differences in total LEC numbers. Notably, similar findings were observed in the iliac LN (**Supplemental Figure 5B-D**) although the magnitude of the effects was reduced compared to the popliteal LN, suggesting that the detrimental effects of WT CHIKV infection on LN structure and function are reduced as cells and antigen move farther downstream in the lymphatic drainage network. Overall, these data suggest that pathogenic CHIKV infection impairs the ability of LECs to acquire antigen upon a secondary foreign challenge, which could have implications for the strength and success of downstream adaptive immunity to respond to that challenge.

**Figure 7.**
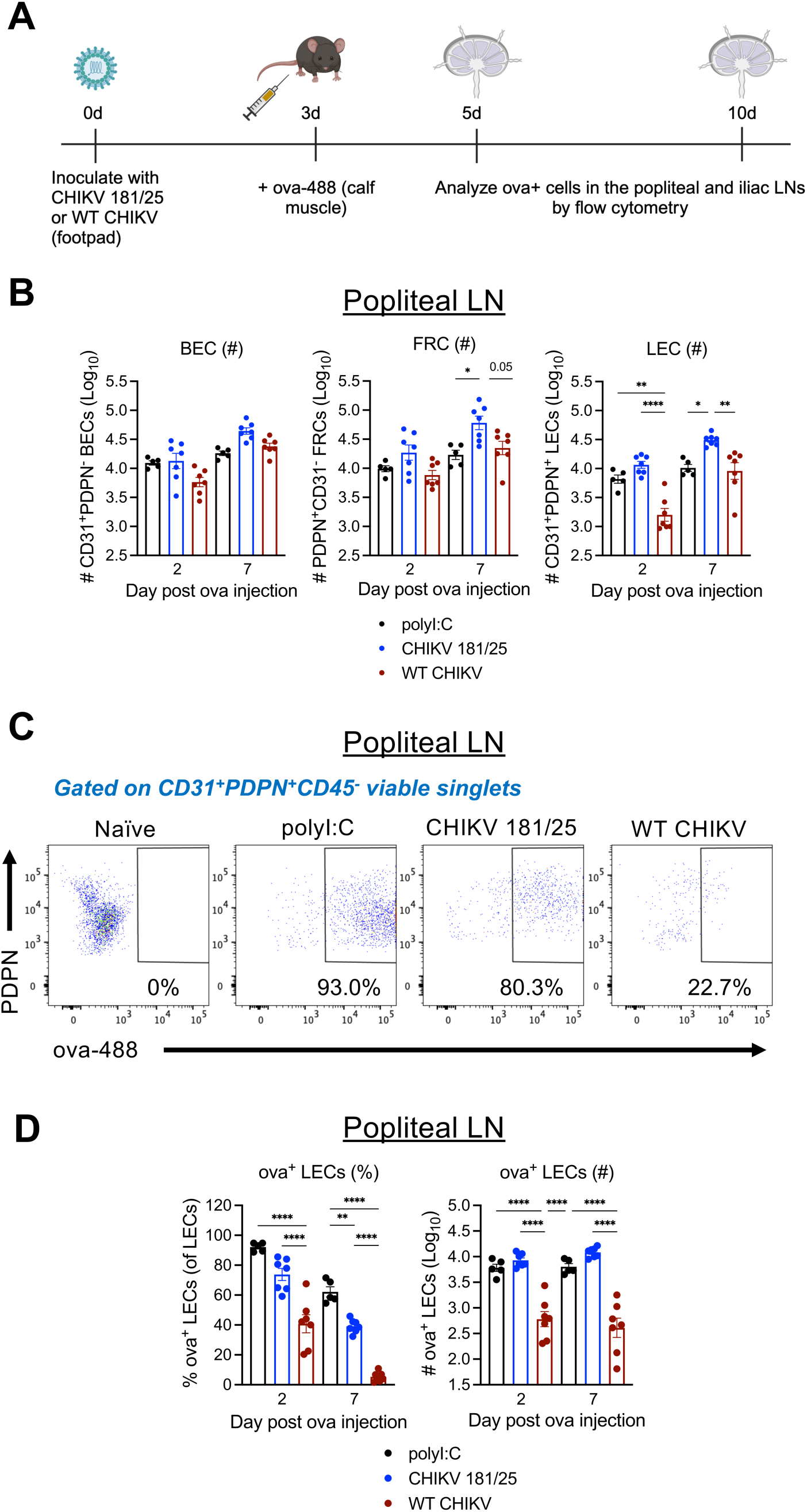
Antigen acquisition by LECs is impaired during WT CHIKV infection. (**A**) WT C57BL/6 mice were mock-inoculated (n = 5) or inoculated in the footpad with 10^3^ PFU CHIKV 181/25 (n = 8) or WT CHIKV (n = 8). At 72 h post-infection, mice were inoculated with 10 µg ova-488 in both calf muscles (20 µg total). As a positive control, naïve mice were injected with 10 µg ova-488 and 5 µg polyI:C in both calf muscles. Ova^+^ LECs in the popliteal LN were then enumerated by flow cytometry at the indicated timepoints. (**B**) LNSC numbers in the popliteal LN following ova immunization. (**C**) Representative flow cytometry plots showing ova^+^ LECs. (**D**) The percentage and total number of ova^+^ LECs. Data are combined from of 2 independent experiments. Statistical significance was determined by one-way ANOVA with Tukey’s multiple comparison test. *, *P* <0.05; **, *P* < 0.01; ***, *P* < 0.001; ****, *P* < 0.0001.

## DISCUSSION

Previous work demonstrated that LN cellular organization and immune responses are disrupted during CHIKV infection (19, 22), however, the specific virus-host interactions that promote these aberrant LN responses remained unknown. Recently, we found that CHIKV particles are internalized by MARCO^+^ medullary LECs in the LN (41). Building on these findings, here we show that throughout the first 24 h of infection, CHIKV RNA accumulates within the floor and medullary subsets of LN LECs, that this accumulation is associated with expression of the scavenger receptor MARCO by these cells, and that these cells may support active viral replication. Moreover, CHIKV infection was associated with the rapid induction of an antiviral and inflammatory gene expression program across LNSCs that was accelerated by CHIKV-MARCO interactions. In addition, we found that as CHIKV infection progressed, both floor and medullary LECs diminished in number in a MARCO-dependent manner. This unique viral targeting and/or capture by LN LECs raised questions about how virus interactions with LNSCs influence the function of these cells in immunity. Indeed, we found that acquisition of vaccine antigen by LECs was reduced following pathogenic CHIKV infection, suggesting that virus-LNSC interactions can influence subsequent secondary responses.

Both previous work and our data here indicate that CHIKV RNA accumulates within specific subsets of LECs in the dLN (41). In both cases, CHIKV RNA preferentially accumulated in LECs expressing higher levels of the scavenger receptor MARCO (41), emphasizing the critical role of MARCO in facilitating CHIKV targeting of LECs rather than indicating a preference for a regionally distinct LEC subset. These data are consistent with our recent report that MARCO can facilitate viral internalization into cells *in vitro* (73) and suggest LECs may be generally less susceptible to CHIKV entry apart from specific subsets. The accumulation of CHIKV RNA in MARCO^+^ floor LECs and medullary MARCO^+^ LECs also reflects the transit over time of viral particles through the LN sinuses. Our previous work has shown CHIKV internalization by these LECs occurs in a MARCO-dependent manner (41), prompting the question of whether these cells are permissive to viral replication. Here, we identified an increased ratio of CHIKV sgRNA within MARCO^+^ LECs at 24 h post-infection and both a lower number of expressed host genes and a higher percentage of mitochondrial reads in those cells, suggesting active viral replication. These data contrast with a prior report that indicated LNSCs are not targets of CHIKV infection using flow cytometry (22). One explanation could be that there is some viral RNA replication within these LECs but the infection is ultimately abortive due to little to no virion production and release. Indeed, expression of the antiviral factor *Bst2*/tetherin was increased at both 8 and 24 h post-infection in LECs, and we observed robust expression of type I IFN-stimulated genes at 24 h post-infection, which together could limit virus replication. Alternatively, due to the small population of LECs that interact with CHIKV, both CHIKV^+^ cell number and the amount of virus production may be too low to detect by methods less sensitive than RNA sequencing. Although CHIKV can interact with a multitude of cell surface proteins for attachment on many cell types, including endothelial cells, other studies have predominantly identified fibroblasts, skeletal muscle cells, and macrophages as targets for active CHIKV replication (15, 16, 57, 58, 73, 75, 76). Indeed, signs of active viral infection in LNSCs are unique and have rarely been identified. Instead, much research has focused on LN resident macrophages, which interact with and can be targeted by viruses such as Zika virus, vaccinia virus, and vesicular stomatitis virus (77–79). However, lymphocytic choriomeningitis virus (LCMV) has been shown to target murine FRCs within lymphoid organs, and importantly, LCMV infection of FRCs was significantly higher for the chronic strain clone 13, which establishes a persistent infection, than the acute strain Armstrong, which is cleared efficiently by CD8^+^ T cells (80). In contrast, while multiple studies have shown that FDCs can trap HIV-1 and maintain infectious virions, representing a potential viral reservoir during chronic infection, FDCs are not permissive to HIV-1 replication and thus viral accumulation in FDCs appears to be an unintended consequence rather than the result of viral targeting of FDCs (81, 82). While more investigation is needed, viral targeting of LNSCs by CHIKV and LCMV, two viruses associated with chronic disease, suggest that viral targeting of LNSCs may have a critical role in the establishment of chronic viral infection. Ultimately, further work is needed to definitively understand the propensity of endothelial cell subsets to support specific stages of the CHIKV infection cycle, including virion internalization, replication, sgRNA translation, and virion assembly and egress.

A key marker of the floor and medullary LEC populations that harbor CHIKV is Lyve-1, which distinguishes them from other LEC subsets, including ceiling, collecting, and valve LECs (24, 25, 29, 40, 63). LNs from WT CHIKV but not attenuated CHIKV 181/25-infected mice displayed a reduced and spatially altered expression of Lyve-1 by 48 h post-infection as well as a complete loss of MARCO expression, suggesting that WT CHIKV infection disrupts Lyve-1^+^ LECs. In lymphatic vessels, Lyve-1 expression is negatively regulated by inflammatory cytokines such as IFN-*γ* and TNF-α (83, 84). Notably, WT CHIKV infection of *MARCO^-/-^* mice did not disrupt Lyve-1 expression, suggesting that the perturbations of Lyve-1 expression observed in CHIKV-infected WT mice could be driven by MARCO-promoted inflammatory responses. Consistent with this idea, we found that the presence of MARCO accelerated and promoted inflammatory gene expression in the LN. Furthermore, we observed numerous CD11b^+^ cells within the disrupted LN sinuses at 48 h post-infection, and the presence of MARCO contributed to the accumulation of inflammatory monocytes in the dLN. Additional work is ongoing to discern if the perturbations of Lyve-1 expression and/or Lyve-1^+^ LECs is immune cell-mediated or LEC-intrinsic and how MARCO influences specific inflammatory factors known to regulate Lyve-1 expression. Interestingly, in one study which evaluated LN LEC subsets following stimulation with imiquimod, a TLR7 agonist, expression of Lyve-1, CD36, and MARCO in floor, ceiling, and medullary LECs remained unaltered (63), in contrast to our findings during CHIKV infection, which also can activate TLR7 (85, 86), suggesting that the altered expression of these molecules during CHIKV infection is more complex.

Despite the altered expression and spatial distribution of Lyve-1 expression observed by confocal microscopy, total LEC, medullary, and floor LEC numbers remained similar in LNs from mock-, CHIKV 181/25-, and WT CHIKV-infected mice at 48 h post-infection when evaluated by flow cytometry. However, by 5 d post-infection, we detected decreased numbers of medullary and floor LECs in LNs from WT CHIKV-infected mice compared to those from mice infected with CHIKV 181/25, which induces less LN inflammation and does not disrupt LN cellular organization (19, 22). These changes were specific to floor and medullary LECs, the LEC subsets that harbor CHIKV RNA based on scRNA-seq. Notably, both floor and medullary LEC numbers were restored in LNs from WT CHIKV-infected *MARCO^-/-^* mice, suggesting that these changes are a consequence of CHIKV targeting of MARCO-expressing LECs. Total LEC numbers remained constant over time during WT CHIKV infection in contrast to CHIKV 181/25 infection; it is possible that this failure of LECs to expand during WT CHIKV infection could be related to alterations in the Prox1-regulated cell differentiation pathway controlling commitment to the LEC or BEC lineages (87–89). Unexpectedly, we observed a reduced proportion of ceiling LECs among total LECs during CHIKV 181/25 infection compared with WT CHIKV at 5 d post-infection, yet total ceiling LEC numbers were similar. It is possible that both *Cd36* and *Icam1* expression are altered in ceiling LECs because of the various immune cell-activating cytokines and chemokines expressed in the dLN in the first 5 d post-infection, biasing our gating analysis. Furthermore, we also observed a reduced proportion of ceiling LECs in WT CHIKV-infected *MARCO^-/-^* mice similar to that seen in CHIKV 181/25-infected WT mice, supporting the idea that this change is part of a functional LN immune response. One caveat to comparison of our flow cytometry and confocal microscopy data is that the major LEC subsets were identified using different surface proteins, as LYVE-1 and MARCO antibodies perform poorly for flow cytometric analysis. Thus, there could be microenvironment-dependent changes in expression of key markers of LEC subsets that complicate interpretation of the cell populations identified by flow cytometry. Indeed, upregulation of ITGA2B has been observed on LECs in response to inflammation (60, 90). Notably, comparison of LEC subsets during homeostatic and response-to-stimuli conditions using scRNA-seq suggested that floor LECs undergo the greatest transcriptional alteration following stimulus with the TLR7 agonist imiquimod, with moderate alteration observed in medullary LECs, and the least changes observed in ceiling LECs (63). However, we note that total LEC numbers in the dLN, as determined by CD31 and PDPN, were reduced during WT CHIKV infection compared with CHIKV 181/25 infection, suggesting that the major changes in LEC subsets are accurate. A previous report investigating the mechanisms dictating expansion and contraction of LN LECs during an immune response demonstrated that type I IFN and PD-L1 both inhibit early LEC division and that a decrease in PD-L1 was sufficient to increase LEC proliferation (91). Given the inflammatory response in the dLN that occurs during WT CHIKV infection, including upregulation of PD-L1 at 24 h post-infection (19, 91), signaling by type I IFN-stimulated genes, such as PD-L1, may prevent expansion of LN floor LECs.

As previously stated, LEC functions include maintenance of LN homeostatic chemokine gradients, guidance to migrating dendritic cells into the LN parenchyma, acquisition and storage of antigen to promote memory CD8^+^ T cell responses, and maintenance of peripheral self-tolerance by AIRE-independent antigen presentation on MHC-II (33–40, 42). Antigen acquisition and retention by LECs supports stronger memory CD8^+^ T cell responses through antigen exchange with, and presentation by, migratory DCs (38, 39). We found that the capacity of LN LECs to acquire antigen after a secondary immunization was specifically impaired during WT CHIKV infection. In contrast, LEC antigen acquisition in mice infected with CHIKV 181/25 was more similar to that observed in naïve mice stimulated with polyI:C, consistent with published data showing that viral infection can also induce LEC antigen acquisition (38). Antigen acquisition was markedly reduced 2 days after immunization in WT CHIKV infected mice, and retention of that antigen also may be impaired as the number of ova^+^ LECs remained constant between 2- and 7-days post-immunization in CHIKV 181/25-infected mice but decreased in WT CHIKV-infected mice. Overall, these data provide evidence that LNSC-targeting viruses that disrupt the function of the cells represent a challenge for vaccination campaigns as patients recently infected with such a virus may need to delay immunization to generate stronger, more protective vaccine-specific responses.

In summary, our work demonstrates that CHIKV interactions with specific subsets of LECs expressing the scavenger receptor MARCO is associated with remodeling of the LNSC transcriptome, extensive LN inflammation, and dysfunction of LN LECs. These findings suggest CHIKV-LEC interactions contribute to impaired downstream LN function and impaired adaptive immunity during CHIKV infection.

## MATERIALS AND METHODS

### Cells

BHK-21 cells (ATCC CCL-10) were grown at 37°C and 5% CO2 in α-minimal essential media (Life Technologies) supplemented with 10% fetal bovine serum (HyClone), 10% tryptose phosphate broth, penicillin and streptomycin, and 0.29 mg/mL L-glutamine.

### Viruses

CHIKV AF15561 (WT CHIKV) is an Asian genotype strain isolated from a human patient in Thailand (GenBank accession no. EF452493). CHIKV AF15561, CHIKV AF15561^E2^ ^K200R^ and CHIKV 181/25 were generated by electroporation of cDNA clone-derived RNA into BHK-21 cells as previously described (92). Briefly, plasmids were linearized by NotI digestion and used as a template for *in vitro* transcription with SP6 DNA-dependent RNA polymerase (Ambion). RNA transcripts were electroporated into BHK-21 cells and at 24 h post-electroporation, cell culture supernatant was collected and clarified by centrifugation at 1,721 x *g*. Clarified supernatants were aliquoted and stored at −80°C. Viral titers were determined by plaque assay using BHK-21 cells, by quantification of RNase-resistant viral genomes by RT-qPCR, or by focus formation assay on Vero cells as previously described (13, 22, 92).

### Mouse Experiments

Mice were bred in specific-pathogen-free facilities at the University of Colorado Anschutz Medical Campus. All mouse studies were performed in an animal biosafety level 3 laboratory. WT C57BL/6J (Jax# 000664) mice were acquired from Jackson Laboratories, and congenic *MARCO^-/-^* mice were provided by Dawn Bowdish (McMaster University) (49). WT and *MARCO^-/-^* mice were housed and bred in specific pathogen-free facilities at the University of Colorado Anschutz Medical Campus. Mice were anesthetized with isoflurane vapors and inoculated with the indicated dose of virus or virus particles in a 10 μl volume via subcutaneous (s.c.) injection into the rear footpad. For scRNA-seq and flow cytometry experiments, mice were inoculated with an equal dose of virus in both rear footpads; for microscopy, mice were inoculated in a single footpad. As sex-based difference have not been observed in the CHIKV infection model, WT male mice were purchased commercially and were age matched and distributed randomly across groups. Based on prior studies of lymph node inflammation during CHIKV infection, mice 4 weeks of age were used in all experiments (19, 22). Experimental animals were humanely euthanized at defined endpoints by exposure to isoflurane vapors followed by bilateral thoracotomy.

### Preparation of single-cell suspensions for single-cell mRNA sequencing

The draining popliteal lymph node from mock- or CHIKV-inoculated mice was pooled into individual replicates (3 replicates; LNs from 5 mice pooled per replicate). Lymph nodes were mechanically homogenized using a 22G needle in Click’s medium (Irvine Scientific, 9195) supplemented with 5 mg/mL liberase DL (Roche, 05401160001) and 2.5 mg/mL DNase (Roche 10104159001) for 1 h at 37°C. After incubation, digested LNs were clarified by passing through a 100-μm cell strainer. Cell suspensions were enriched for CD45^-^ cells by labeling cells with PE-conjugated anti-mouse CD45 (30-F11), CD140A (APA5), and Ter119 (Ter119) monoclonal antibodies and subsequent depletion of PE-labeled cells using Miltenyi anti-PE microbeads (130-048-801) and Miltenyi MACS LS (130-042-401) columns according to the manufacturer’s instructions with the following modifications: (i) We used 25% of the recommended volume of anti-PE microbeads, and (ii) we subjected the CD45^-^ enriched cell fraction to a second MACS LS column. All cell suspensions post-column enrichment were enumerated using a hemacytometer. Cell fractions throughout the procedure were analyzed for PE-labeled cell depletion and enrichment of CD45^-^ cells by flow cytometry. Cell fractions were stained with fixable LIVE/DEAD dye (Invitrogen, L34955) and antibodies against the following cell surface antigens: CD45 (30-F11), CD31 (390), PDPN (8.1.1), B220 (RA3-6B2), TCRβ (H57-597), CD11b (M1/70), and Ly6C (HK1.4). All flow cytometry antibodies were obtained from BioLegend, BD Bioscience, or eBioscience. Following surface antigen staining, cells were washed and fixed in 1% paraformaldehyde (PFA)/1% FBS, and data were acquired on a BD LSR Fortessa X-20 flow cytometer. Data analysis was performed using FlowJo analysis software (Tree Star).

### Single-cell library preparation using the 10× Genomics platform

LN cell suspensions enriched for CD45^-^ cells were subjected to single-cell droplet-encapsulation using the Next GEM Chip G Kit (1000127) and a 10× Genomics chromium controller housed in a BSL3 laboratory. We targeted recovery of 10,000 cells for single-cell RNA sequencing for each replicate. Single-cell gene expression libraries were generated using the Next GEM single-cell 30 GEM library and gel bead kit v3.1 (1000128) and single index kit T set A (1000213) according to the manufacturer’s instructions (10× Genomics). Sequences were generated with the Illumina NovaSEQ 6000 instrument using S4 flow cells and 300 cycle SBS reagents. We targeted 50,000 reads per cell, with sequencing parameters of Read 1:151 cycles; i7 index: 10 cycles; i5 index: 0 cycles; Read 2: 151 cycles.

### CHIKV-capture library preparation and analysis

The scRNA-seq libraries for the 8 h timepoint were enriched for molecules aligning to the CHIKV genome according to a previously published method (55, 93). Specifically, the CHIKV genome was PCR amplified in 3 fragments (primer sequences: CHIKV-F1 5ʹ – TGAGACACACGTAGCCTACCA – 3ʹ, CHIKV-F2 5ʹ-AAGTCCAAGGGAATACAGATCTTC – 3ʹ, CHIKV-F3 5ʹ – ACCGCAGCACGGTAGAGA – 3ʹ, CHIKV-R1 5ʹ – CGAATAACATTACCTTGGAGCA – 3ʹ, CHIKV-R2 5ʹ –TTTTTCCCGGCCTATCACAG – 3ʹ, CHIKV-R3 5ʹ – AAAAACAAAATAACATCTCCTACGTC – 3ʹ) and labeled with biotin-dUTP using the same primers before sonicating to generate ∼150 bp fragments for hybridization. Denatured and diluted biotin-dUTP-labeled CHIKV genome fragments were hybridized to the concentrated scRNA-seq libraries separately. Streptavidin capture beads were mixed with the hybridized libraries and washed to remove unbound DNA. Libraries were amplified directly from the cleaned-up beads and sequenced as described above under **Single-cell library preparation using the 10× Genomics platform**. FASTQ files for each replicate (2 mock- and CHIKV-infected for 8 h post-infection) were processed using the cellranger count pipeline (v5.0.1). Reads were aligned to the mm10 and CHIKV AF15561 (EF452493.1) reference genomes.

Our CHIKV-capture protocol resulted in the identification of ∼4 fold more cells with detectable CHIKV RNA when compared with the unenriched libraries. To quantify overall CHIKV RNA levels for each cell, we calculated a CHIKV score (**Figure 1A and C**), which is the number of CHIKV-capture counts aligning to the CHIKV genome divided by the total CHIKV-capture counts for the cell (mouse counts + CHIKV counts). To visualize this metric on UMAP projections (**Figure 1A**), a pseudo count (smallest non-zero value / 2) was added to each value plotted.

### Single-cell RNA-seq gene expression analysis

FASTQ files for each replicate (2 mock- and CHIKV-infected for 8 h post-infection; 3 mock- and CHIKV-infected for 24 h post-infection) were processed using the cellranger count pipeline (v5.0.1). Reads were aligned to the mm10 and CHIKV AF15561 (EF452493.1) reference genomes. The CHIKV genome included annotations for the sgRNA (position 7567-12036) and 5ʹ (position 1-7566) regions. Initial filtering of gene expression data was performed separately for the 8 h and 24 h timepoints using the Seurat R package (v4.2.0). Gene expression data for each biological replicate were combined into a single Seurat object. CHIKV counts were excluded from the gene expression matrices so they would not influence downstream processing (dimensionality reduction, clustering) of the mouse expression data.

Previously published scRNA-seq data for the 24 h timepoint was processed as previously described (41). CHIKV-low and -high cells were identified by first filtering cells to only include those with >5 CHIKV counts. K-means clustering was then used to independently group each biological replicate into CHIKV-low and -high populations. Cells with 5 CHIKV counts or less were included in the CHIKV-low population. Cells were filtered based on the number of detected mouse genes (>250 and <6000) and the percent mitochondrial counts (<20%). Genes were filtered to only include those detected in >5 cells. Potential cell doublets were removed using the DoubletFinder (v2.0.3) R package using an estimated doublet rate of 10%. Due to the ability of CHIKV to inhibit host transcription, CHIKV-high cells with a low number of detected mouse genes (<250) or a high fraction of mitochondrial reads (>20%) were not filtered and remained in the dataset for the downstream analysis. The sgRNA ratio (**Figure S2B, D, E**) was calculated for the 24 h timepoint by dividing sgRNA (position 7567-12036) counts by 5ʹ (position 1-7566) counts. To visualize this metric on UMAP projections (**Figure S2B**), before calculating the sgRNA ratio, a pseudo count of 1 was added to the sgRNA and 5’ counts for each cell plotted (to eliminate division by 0).

Cells from the 8 h timepoint samples were filtered based on the number of detected mouse genes (>250 and <8000) and percent mitochondrial counts (<20%). Genes were filtered to only include those detected in >5 cells. The cell calls made by the cellranger pipeline (10X Genomics) for the 8 h CHIKV replicate 2 sample were not accurate based on analysis of UMI counts and likely included a substantial number of empty droplets. To account for this, a cutoff of 800 UMI counts was used to remove potential empty droplets from this sample. Counts from the CHIKV-capture libraries were then added to the object for all cells passing our filtering cutoffs. Due to the very few cells with detectable CHIKV RNA at this early 8 h timepoint, CHIKV^+^ cells were classified as any cell with at least 1 CHIKV-capture count aligning to the CHIKV genome.

For both the 8 h and 24 h timepoints, mouse gene expression counts were normalized by the total mouse counts for the cell, multiplied by a scale factor (10,000), and log-transformed (NormalizeData), Normalized mouse counts were scaled and centered (ScaleData) using the top 2000 variable features (FindVariableFeatures). The scaled data were used for PCA (RunPCA) and the first 40 principal components were used to identify clusters (FindNeighbors, FindClusters) and calculate uniform manifold approximation and projection (UMAP) (RunUMAP).

To ensure accurate and consistent cell type annotations, we integrated the 8 h and 24 h datasets based on timepoint (8 h and 24 h) and sample (mock- and CHIKV-infected) using the R package, Harmony (v0.1.1) (94). We then re-clustered the cells using the integrated data and generated an initial set of broad cell type annotations using the R package, clustifyr (v1.8.0) (95) and reference data from Immgen (96). These annotations were checked for accuracy and further refined using known cell type markers including, Cd19 (B cells), Cd3e (T cells), Hba-a1 (erythrocytes), Pdpn, and Pecam1. To identify perivascular cells (PvCs), fibroblasts were re-clustered, integrated, and clusters were annotated using published reference data (Rodda et al., 2018) (as described above). To identify endothelial cell subsets, endothelial cells were re-clustered, integrated, and clusters were annotated using published reference data (Xiang et al., 2020) (as described above). LEC and BEC annotations were further refined using known marker genes including Marco, Pdpn, and Pecam1. Visualization of integrated UMAP projections suggest that broad LN cell type and endothelial cell type annotations were consistent across conditions (**Supplemental Figure 3A and B**). In addition, strong correlation with the published reference data (**Supplemental Figure 3C**) and expression of key endothelial cell marker genes (**Supplemental Figure 3D**) further support the accuracy of our cell type annotations.

Differentially expressed genes were identified for each cell type for mock vs 8 h and 8 h vs 24 h using the Seurat package. To allow for equal comparison with the 8 h timepoint, the top two replicates (based on cell number) from the 24 h timepoint were used for identifying differentially expressed genes. Genes were considered upregulated if the average log2 fold change was >0.15 for 8 h and >0.25 for 24 h for all replicates and the largest p-value for all replicates was <0.05. Gene ontology terms (Biological Process) were identified for the top 200 upregulated genes (sorted by maximum p-value for replicates) for each cell type using the R package, clusterProfiler (v4.4.4) (97). Terms were filtered to only include those with an adjusted p-value <0.05 and at least 3 or 10 upregulated genes overlapping the term for the 8 h and 24 h timepoints, respectively. Terms with <10 or >750 genes were excluded from the analysis. Terms identified for each cell type were combined and clustered into 5 modules based on the pairwise overlap between terms using the clusterProfiler package. Enrichment scores were calculated by dividing the fraction of upregulated genes overlapping the term by the fraction of background genes overlapping the term. For **Figure 5C and E**, cell types are only shown if they have at least one upregulated gene overlapping any of the terms plotted.

### Immunofluorescence and confocal microscopy

LNs were removed at the indicated time post-infection, fixed in 1 mL of periodate-lysine-paraformaldehyde (PLP) buffer containing

0.1 M L-lysine, 2% PFA, and 2.1 mg/mL NaIO4 at pH 7.4 for 24-48 h at 4°C, followed by incubation for at least 24 h in 30% sucrose phosphate-buffered solution. Tissues were embedded in optimal-cutting-temperature (OCT) medium (Electron Microscopy Sciences), carefully oriented to allow for sectioning through both the B cell follicles and the medullary sinuses, and frozen in dry-ice-cooled isopentane. 16-µm sections were cut on a Leica cryostat (Leica Microsystems). Sections were blocked with 5% goat, donkey, bovine, rat, or rabbit serum and then stained with one or more of the following Abs: ERTR-7 (rat monoclonal clone ERTR7, Abcam), B220 (clone RA3-6B2, Thermo Fisher Scientific), Lyve-1 (clone ALY7, Thermo Fisher Scientific), CD11b (clone M1/70, Biolegend), CD36 (HM36), and/or MARCO (clone MCA1849, Serotec). Sections were incubated with secondary antibodies as needed and as controls, and images were acquired using identical PMT (photomultiplier tube) and laser settings. Images were acquired on a Leica SP8 or Stellaris confocal microscope (Leica Microsystems) using a 40X 1.3 NA or 63X 1.4 NA objective and merged to cover the entire LN using the Leica tilescan function. Images were processed and analyzed using Imaris software 8.0 (Oxford Instruments).

### Isolation of cells from LNs and flow cytometry

To evaluate the cellular composition of LNs, nodes were gently homogenized in a Biomasher II tissue homogenizer (Kimble Chase) in RPMI 1640 (HyClone) supplemented with 5% FBS. For LN stromal cell isolation, left and right popliteal LNs from two mice were combined for each sample (4 LNs total), minced in Click’s media (Sigma-Aldrich) with 22G needles (Exelint), and digested for 1 h at 37°C in 94 µg/mL DNase I (Roche) and either 250 µg/mL Liberase DL (Sigma-Aldrich) or 250 µg/mL collagenase type I and 250 µg/mL collagenase type IV (Worthington Biochemicals). LN suspensions were supplemented with Click’s media with 2.5% FBS and 5 mM EDTA. For both stromal cell isolation and general LN cell composition, cell suspensions were passed through a 100-µm cell strainer (BD Falcon) and total viable cells were enumerated by trypan blue exclusion. Single-cell suspensions were first incubated for 15 min at 25 °C in LIVE/DEAD Fixable Violet Dead Cell Stain (Invitrogen) to identify viable cells, then incubated for 20 min at 4°C with anti-mouse Fc*γ*RIII/II (2.4G2; BD Pharmingen) and then stained for 45 min at 4°C with the following antibodies from BioLegend diluted in FACS buffer (PBS with 2% FBS): anti-CD45 (30-F11), anti-CD11b (M1/70), anti-CD11c (N418), anti-Ly6C (HK1.4), anti-Ly6G (1A8), anti-CD31 (390), anti-Podoplanin (8.1.1), anti-CD36 (HM36), anti-CD206/MRC1 (C068C2), anti-CD41/ITGA2B (MWReg30), and anti-CD54/ICAM1

(YN1/1.7.4). Cells were washed three times in PBS/2% FBS and then fixed for 15 min in 1x PBS/1% PFA and analyzed on a BD LSR Fortessa cytometer using FACSDiva software. Further analysis was performed using FlowJo software (Tree Star).

### Gene expression analysis by RT-qPCR

LNs were dissected and homogenized in TRIzol reagent (Life Technologies) with a FastPrep-24 Classic homogenizer (MP Biomedicals). RNA was isolated using the PureLink RNA Mini kit (Thermo Fisher) with on-column DNase treatment. Gene expression was quantified by RT-qPCR using available Taqman gene expression assays (Thermo Fisher). Expression of each gene was normalized to 18S and analyzed as fold change over mock samples.

### Evaluation of antigen acquisition by LNSCs

Antigen acquisition by LNSCs was evaluated using fluorescently labeled ovalbumin (ova) as previously described (38). Ovalbumin (Catalog number A5503, Sigma-Aldrich, St. Louis, MO) was decontaminated of lipopolysaccharide using a Triton X-114 detoxification method and tested with Pierce LAL chromogenic endotoxin quantitation kit (catalog number 88282, Thermo Fisher Scientific, Waltham, MA). Ovalbumin was labeled with Alexafluor 488 using Alexafluor 488 succimidyl ester labeling system (Catalog number A20100, Thermo Fisher Scientific, Waltham, MA). Mice were inoculated with 20 µg AF488-labeled ovalbumin total via intramuscular injection into both calf muscle (10 µg per calf), and both popliteal and iliac LNs collected at the indicated timepoint for analysis of ova^+^ LNSCs by flow cytometry. A naïve mouse was used as a negative control for ova^+^ LNSCs. LNSCs were processed for flow cytometry according to the methods described under **Isolation of cells from lymph nodes and flow cytometry.**

### Statistical Analysis

All non-sequencing data were analyzed using GraphPad Prism version 9.4.1 software. Data were evaluated for statistically significant differences using a two-tailed, unpaired t test, and either a one-way or two-way analysis of variance (ANOVA) test followed by Tukey’s multiple comparison test. A *P*-value < 0.05 was considered statistically significant. P-values were indicated as follows: *, *P* < 0.05; **, *P* < 0.01; ***, *P* < 0.001; ****, *P* < 0.0001. All differences not specifically indicated to be significant were not significant (*P* > 0.05).

### Study approval

All animal experiments conducted at the University of Colorado Anschutz Medical Campus were performed with the approval of the Institutional Animal Care and Use Committee (IACUC) of the University of Colorado School of Medicine (Assurance Number: A3269-01) under protocol 00026.

### Data Availability

The single-cell RNA-seq data underlying Figs. 1, S2, S3, and 5 are available at NCBI GEO under accessions GSE174667 and GSE243638. A reproducible analysis pipeline is available at https://github.com/rnabioco/morrison-lnsc. Differentially expressed genes and gene ontology terms for **Figure 5** are shown in **Table S1**. Values for all data points in Figs. S1, 2, 3, 4, S4, S5, 6, and 7 are available in the Supporting Data Values file.

## AUTHOR CONTRIBUTIONS

CJL and TEM conceptualized the study. CJL, RMS, GVR, BJD, MKM and AM conducted experiments and acquired the data. CJL, RMS, GVR, MKM, HDH, and TEM analyzed the data. BAT and TEM provided resources. CJL and TEM wrote the original draft of the manuscript. CJL, RMS, JRH, HDH, BAT, and TEM reviewed and edited the manuscript. JRH, BAT, HDH, and TEM acquired funding. JRH, BAT, and TEM supervised the study.

## ACKNOWLEDGEMENTS

This research was supported by Public Health Service grants R01 AI141436 and AI148144 from the National Institute of Allergy and Infectious Diseases to T.E.M. C.J.L. was supported by an RNA Biosciences Initiative Research Scholar Award and T32 AI052066. R.M.S. is supported by the RNA Bioscience Initiative and T32 AI074491. J.R.H is supported by grants R35 GM119550 and R01 AG071467. B.A.J.T. is supported by NIH R01 AI121209, R21 AI155929 and R01 AI155474. H.D.H is supported by the Intramural Research program of NIAID. This research also was supported by the CU Anschutz SOM Programmatic Incubator for Research (CU ASPIRE) Program. The funders had no role in study design, data collection and interpretation, or the decision to submit the work for publication. Flow cytometry was performed in the University of Colorado Cancer Center Flow Cytometry Shared Resource (RRID:SCR_022035). The authors have no conflicting financial interests.

## SUPPLEMENTAL FIGURE LEGENDS

**Figure S1. Enrichment of CD45^-^ LN stromal cells for scRNA-seq.** (**A-E**) WT C57BL/6 mice were mock-inoculated (n = 2) or inoculated in both rear footpads with 10^3^ PFU CHIKV (n = 2). The left and right popliteal LNs were collected at 8 h post-infection for enrichment of CD45^-^ LN stromal cells (LNSCs) via depletion of CD45^+^ cells. The proportion of CD45^+^ and CD45^-^ cells was evaluated pre- and post-depletion by flow cytometry. (**A**) Representative flow cytometry plots showing the gating strategy for live CD45^-^ LNSCs. (**B and C**) Representative flow cytometry plots of live cell viability (**B**) and percentage of CD45^-^ cells (**C**) in mock and CHIKV-infected samples pre- and post-CD45^+^ cell depletion. (**D and E**) Percentage of CD45^+^ and CD45^-^ cells in each condition and replicate pre-(**D**) and post-(**E**) CD45^+^ cell depletion.

**Figure S2. Signs of CHIKV replication in MARCO-expressing LECs.** (**A-E**) WT C57BL/6 mice were inoculated with PBS (mock, n = 3) or 10^3^ PFU of CHIKV (n = 3) in the left-rear footpad. At 24 h post-infection, the dLN was collected and enzymatically digested into a single-cell suspension. Cells were enriched for CD45^-^ cells and analyzed by scRNA-seq as previously described (41). (**A**) UMAP projection shows annotated cell types. (**B**) UMAP projection shows CHIKV sgRNA ratio (sgRNA counts/5’ counts). (**C**) The fraction of cells identified as CHIKV-high is shown for each cell type. Labels show the number of CHIKV-high cells/total cells. P values were calculated as described in **Figure 1D**. (**D**) CHIKV sgRNA ratio for cells with >0 sgRNA counts and >0 5’ counts. Only cell types with >40 cells are shown. P values were calculated using a two-sided Wilcoxon rank sum test with Bonferroni correction. (**E**) The correlation between CHIKV sgRNA ratio and QC metrics for CHIKV-high MARCO^+^ LECs and unassigned-LECs.

**Figure S3. Cell type annotation of scRNA-seq data.** WT C57BL/6 mice were inoculated with PBS (mock, n = 3) or 10^3^ PFU of CHIKV (n = 3) in the left-rear footpad. At 8 and 24 h post-infection, the dLN was collected and enzymatically digested into a single-cell suspension. Cells were enriched for CD45^-^ cells and analyzed by scRNA-seq as previously described (41). (**A**) UMAP projections of cell type annotations for integrated data. (**B**) UMAP projections of endothelial cell type annotations for integrated data. (**C**) Correlation between annotated LEC subsets and reference data. (**D**) Expression of select marker genes across LN cells.

**Figure S4. Lyve-1 and MARCO expression over time during WT and attenuated CHIKV infection.** (**A-C**) WT C57BL/6 mice were mock-inoculated (n = 3) or inoculated in the footpad with 10^3^ PFU CHIKV 181/25 (n = 5) or WT CHIKV (n = 5). At 8 (**A**), 24 (**B**), or 48 (**C**) h post-infection the dLN was collected. Frozen dLN sections were stained for B220 (B cells; blue), Lyve-1 (LECs; white), and MARCO (red). Scale bar, 200 μm. Images are representative of 3-5 dLNs per group (2 independent experiments).

**Supplemental Figure 5. Impaired antigen acquisition is LEC-specific and not limited to the popliteal LN.** 4 week old WT C57BL/6 mice were mock-infected or infected in the footpad with 10^3^ PFU CHIKV 181/25 or WT CHIKV. At 72 h post-infection, mice were inoculated with 10 µg ova-488 in both calf muscles (20 µg total). As a positive control, naïve mice were injected with 10 µg ova-488 and 5 µg polyI:C in both calf muscles. Ova^+^ LNSCs in the popliteal and iliac LNs were then evaluated by flow cytometry at the indicated timepoints. Representative plots showing ova^+^ LNSCs in the popliteal LN (**A**). Percentage of ova^+^ BECs, FRCs, and LECs among each condition in the popliteal LN. (**B**) LNSC numbers in the iliac LN following ova immunization (**C**). Representative plots showing ova^+^ LECs in the iliac LN, including the naïve control for gating on ova^+^ LECs (**D**) and quantification of percentage and number of ova^+^ LECs (**E**). Data representative of 2 independent experiments. Statistical significance was determined by two-way ANOVA with Tukey’s multiple comparison test. Only comparison of ova^+^ BECs, FRCs, and LECs within each condition is shown. ***, *P* < 0.01; ****, *P* < 0.0001.

