## Supplemental Figures S1-S5 for "Chikungunya virus infection disrupts lymph node lymphatic endothelial cell composition and function via MARCO"

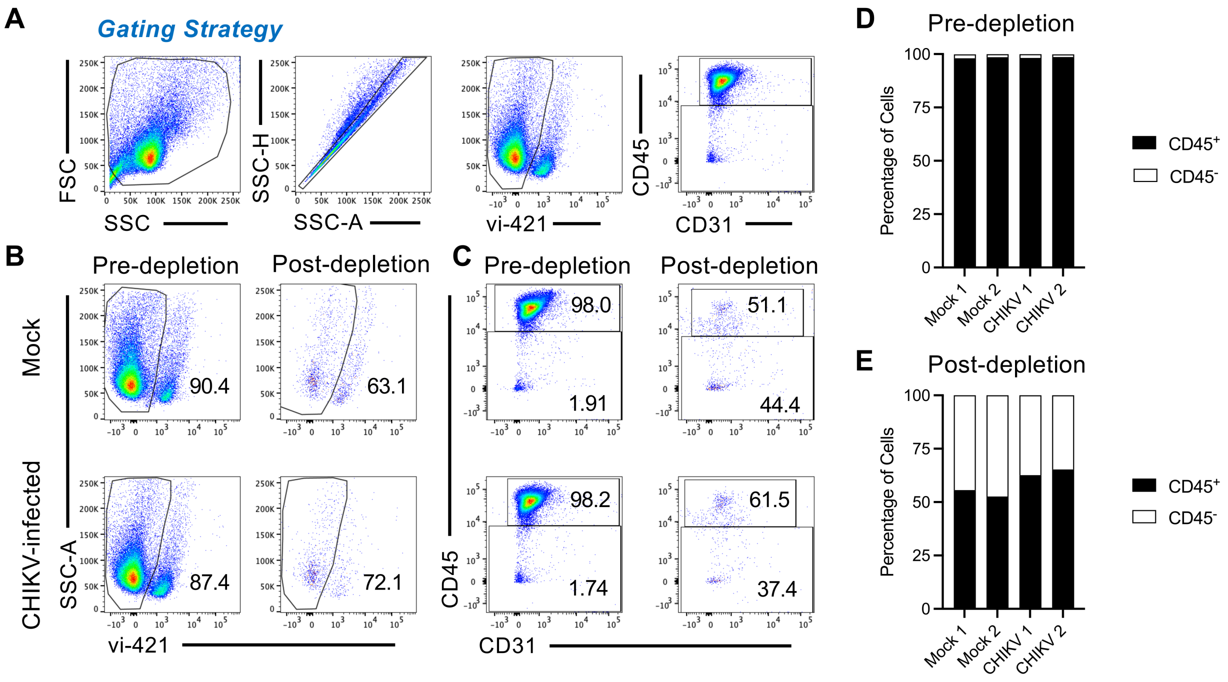


**Figure S1. Enrichment of CD45- LN stromal cells for scRNA-seq.** (**A-E**) WT C57BL/6 mice were mock-inoculated (n = 2) or inoculated in both rear footpads with $10^3$ PFU CHIKV (n = 2). The left and right popliteal LNs were collected at 8 h post-infection for enrichment of CD45- LN stromal cells (LNSCs) via depletion of CD45+ cells. The proportion of CD45+ and CD45- cells was evaluated pre- and post-depletion by flow cytometry. (**A**) Representative flow cytometry plots showing the gating strategy for live CD45- LNSCs. (**B and C**) Representative flow cytometry plots of live cell viability (**B**) and percentage of CD45- cells (**C**) in mock and CHIKV-infected samples pre- and post-CD45+ cell depletion. (**D and E**) Percentage of CD45+ and CD45- cells in each condition and replicate pre- (**D**) and post-(**E**) CD45+ cell depletion.

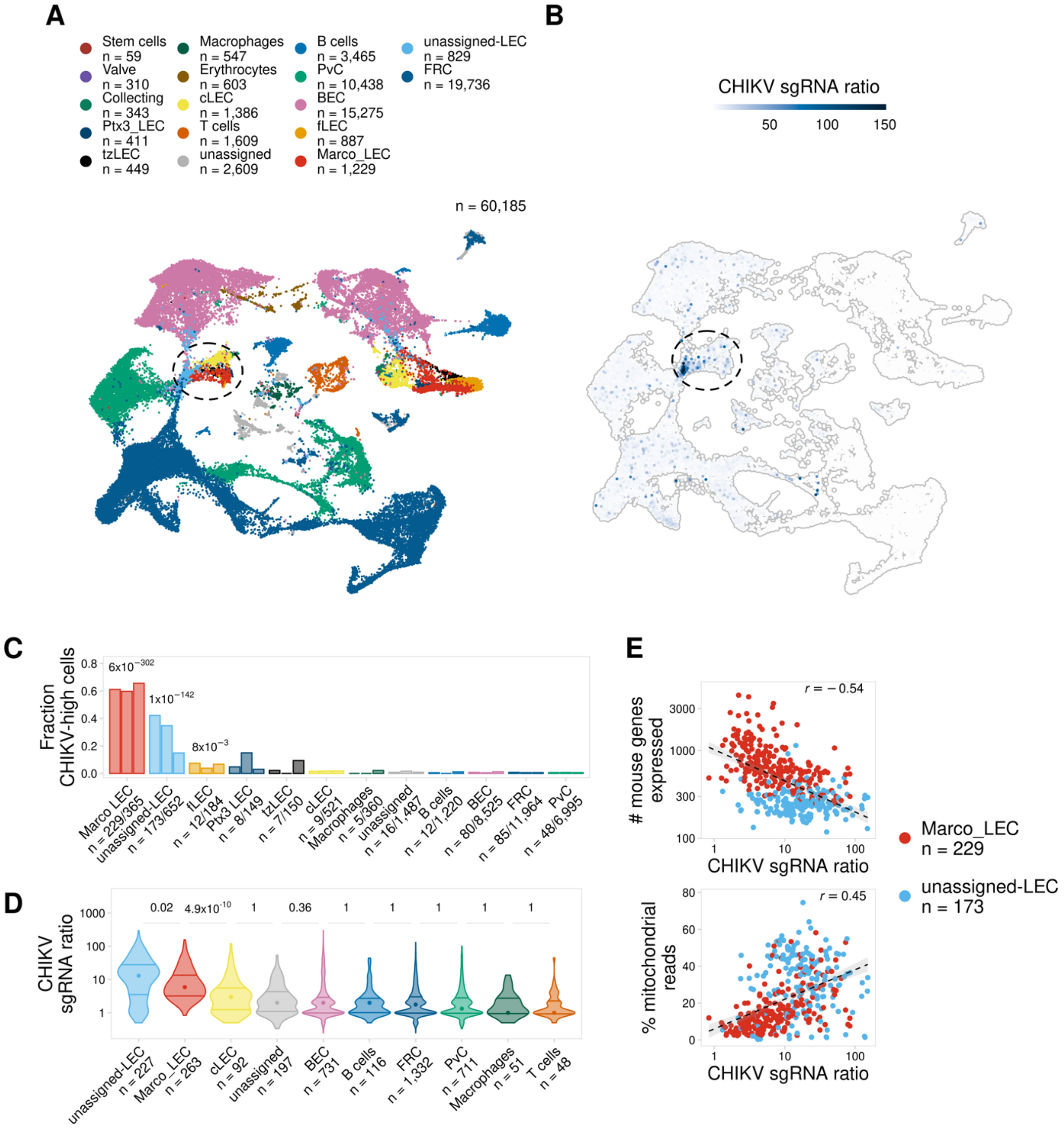


**Figure S2. Signs of CHIKV replication in MARCO-expressing LECs.** (**A-E**) WT C57BL/6 mice were inoculated with PBS (mock, n = 3) or $10^3$ PFU of CHIKV (n = 3) in the left-rear footpad. At 24 h post-infection, the dLN was collected and enzymatically digested into a single-cell suspension. Cells were enriched for $CD45^-$ cells and analyzed by scRNA-seq as previously described (Carpentier et al., 2021). (**A**) UMAP projection shows annotated cell types. (**B**) UMAP projection shows CHIKV sgRNA ratio (sgRNA counts/5' counts). (**C**) The fraction of cells identified as CHIKV-high is shown for each cell type. Labels show the number of CHIKV-high cells/total cells. P values were calculated as described in **Figure 1D**. (**D**) CHIKV sgRNA ratio for cells with >0 sgRNA counts and >0 5' counts. Only cell types with >40 cells are shown. P values were calculated using a two-sided Wilcoxon rank sum test with Bonferroni correction. (**E**) The correlation between CHIKV sgRNA ratio and QC metrics for CHIKV-high MARCO$^+$ LECs and unassigned-LECs.

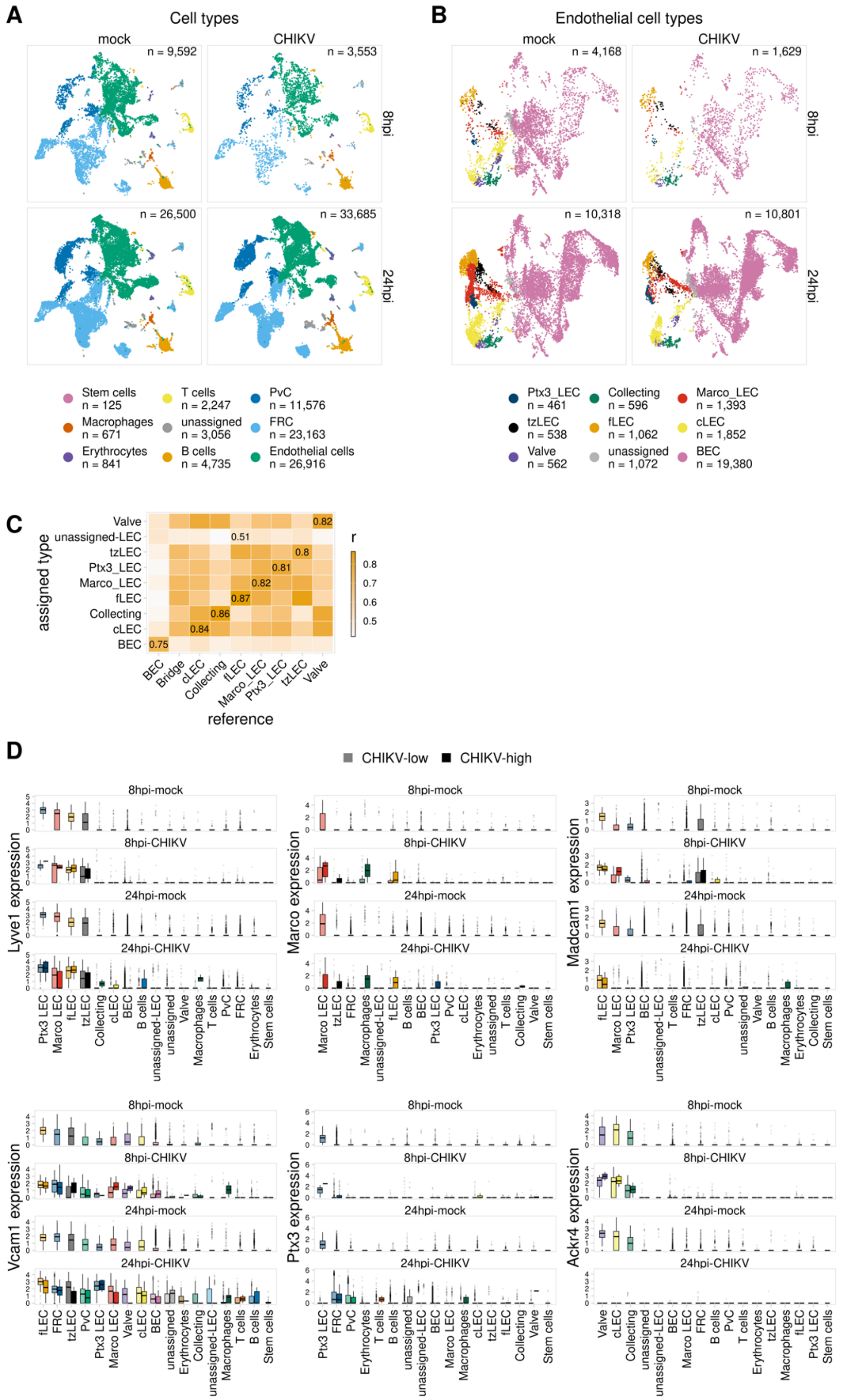


**Figure S3. Cell type annotation of scRNA-seq data.** WT C57BL/6 mice were inoculated with PBS (mock, n = 3) or $10^3$ PFU of CHIKV (n = 3) in the left-rear footpad. At 8 and 24 h post-infection, the dLN was collected and enzymatically digested into a single-cell suspension. Cells were enriched for $CD45^-$ cells and analyzed by scRNA-seq as previously described (Carpentier et al., 2021). (**A**) UMAP projections of cell type annotations for integrated data. (**B**) UMAP projections of endothelial cell type annotations for integrated data. (**C**) Correlation between annotated LEC subsets and reference data. (**D**) Expression of select marker genes across LN cells.

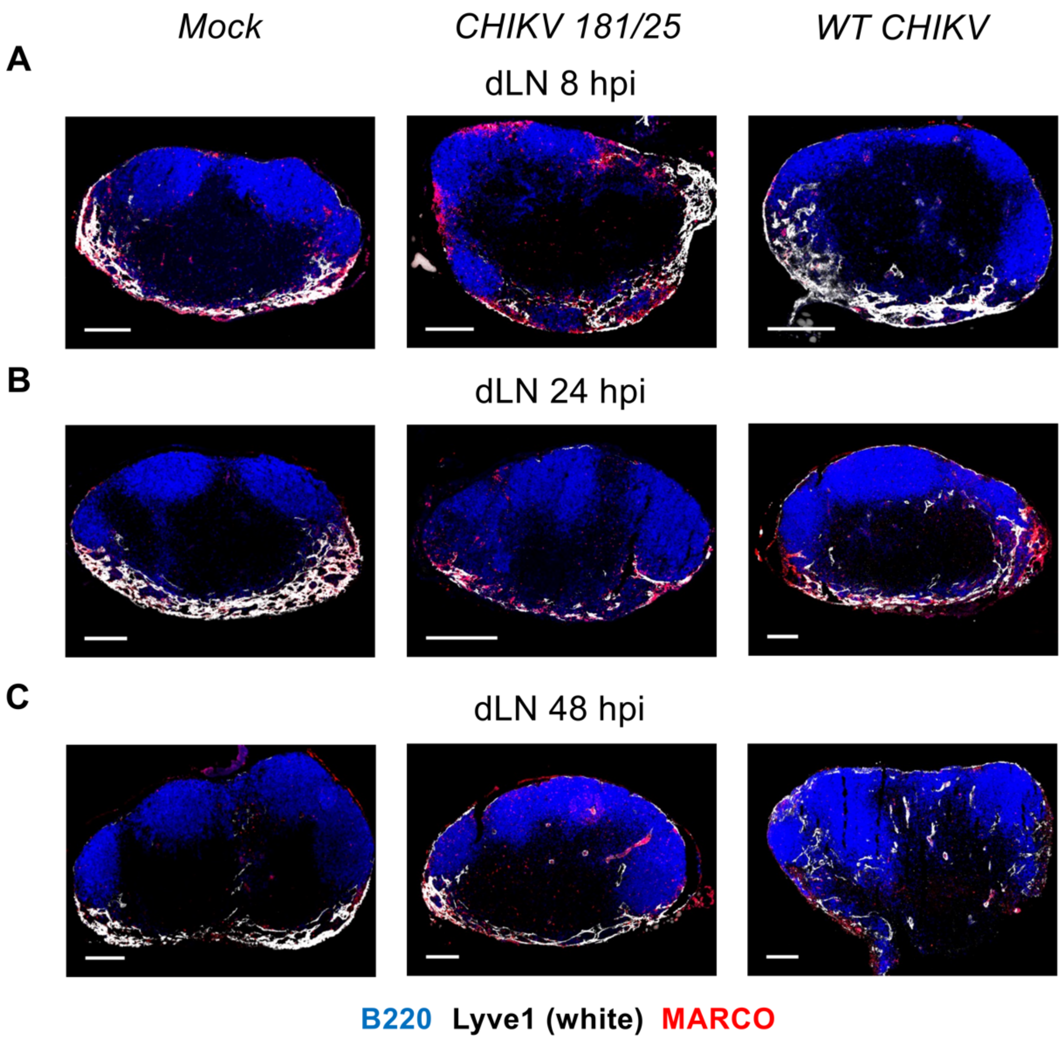


**Figure S4. Lyve-1 and MARCO expression over time during WT and attenuated CHIKV infection.** (**A-C**) WT C57BL/6 mice were mock-inoculated (n = 3) or inoculated in the footpad with $10^3$ PFU CHIKV 181/25 (n = 5) or WT CHIKV (n = 5). At 8 (**A**), 24 (**B**), or 48 (**C**) h post-infection the dLN was collected. Frozen dLN sections were stained for B220 (B cells; blue), Lyve-1 (LECs; white), and MARCO (red). Scale bar, 200 μm. Images are representative of 3-5 dLNs per group (2 independent experiments).

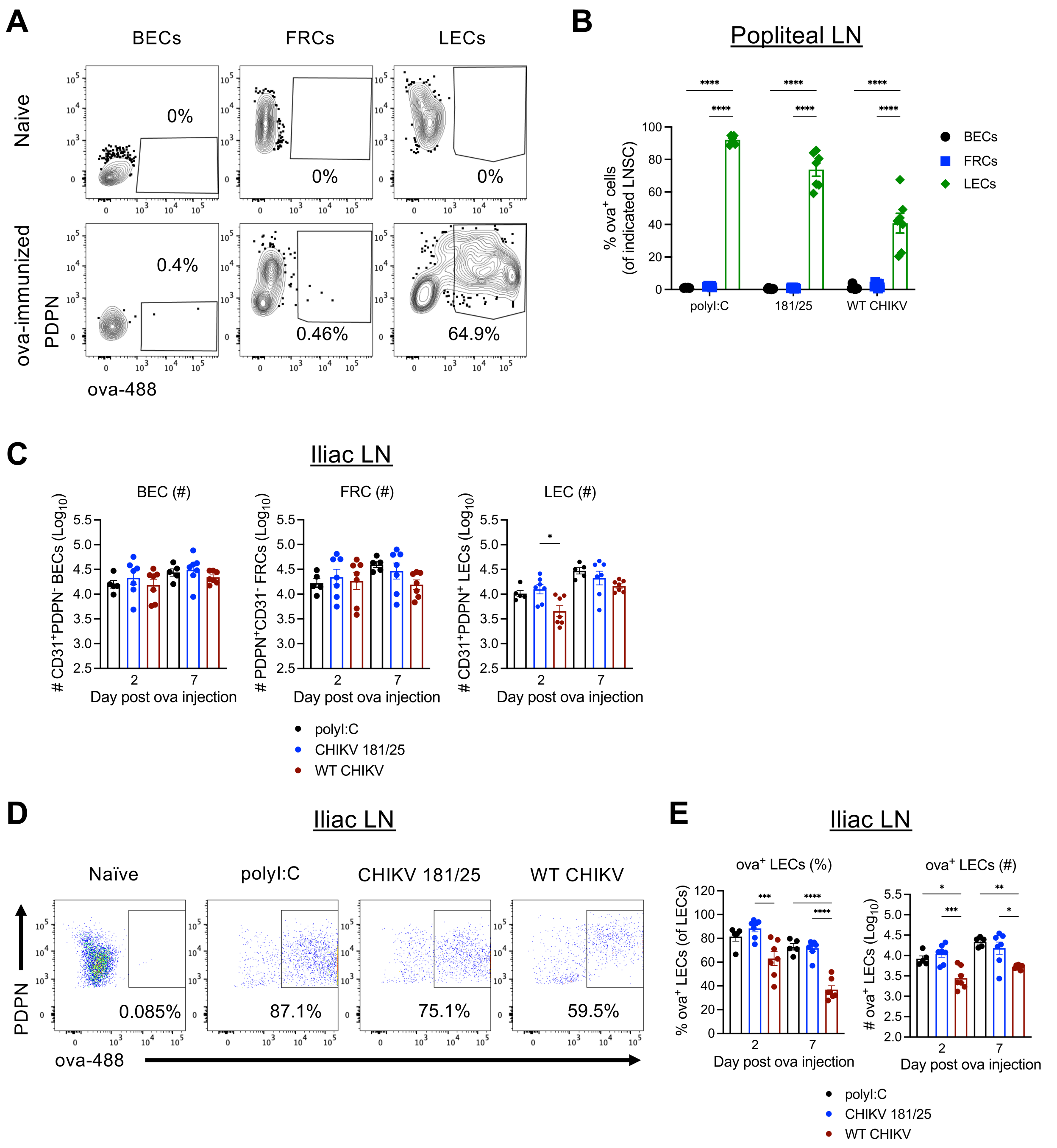


**Supplemental Figure 5. Impaired antigen acquisition is LEC-specific and not limited to the popliteal LN.** 4 week old WT C57BL/6 mice were mock-infected or infected in the footpad with $10^3$ PFU CHIKV 181/25 or WT CHIKV. At 72 h post-infection, mice were inoculated with 10 µg ova-488 in both calf muscles (20 µg total). As a positive control, naïve mice were injected with 10 µg ova-488 and 5 µg polyI:C in both calf muscles. Ova$^+$ LNSCs in the popliteal and iliac LNs were then evaluated by flow cytometry at the indicated timepoints. Representative plots showing ova$^+$ LNSCs in the popliteal LN (**A**). Percentage of ova$^+$ BECs, FRCs, and LECs among each condition in the popliteal LN. (**B**) LNSC numbers in the iliac LN following ova immunization (**C**). Representative plots showing ova$^+$ LECs in the iliac LN, including the naïve control for gating on ova$^+$ LECs (**D**) and quantification of percentage and number of ova$^+$ LECs (**E**). Data representative of 2 independent experiments. Statistical significance was determined by two-way ANOVA with Tukey's multiple comparison test. Only comparison of ova$^+$ BECs, FRCs, and LECs within each condition is shown. ***, $P < 0.01$; ****, $P < 0.0001$.
